# Sublethal pesticide exposure decreases mating and disrupts chemical signaling in a beneficial pollinator

**DOI:** 10.1101/2024.12.04.626868

**Authors:** Nathan Derstine, Cameron Murray, Freddy S Purnell, Etya Amsalem

**Author notes:** Nathan Derstine (corresponding author) –. Cameron Murray -, Frederick Purnell -, Etya Amsalem –.

## Abstract

Pesticides provide vital protection against insect pests and the diseases they vector but are simultaneously implicated in the drastic worldwide decline of beneficial insect populations. Convincing evidence suggests that even sublethal pesticide exposure has detrimental effects on both individual- and colony-level traits, but the mechanisms mediating these effects remained poorly understood. Here, we use bumble bees to examine how sublethal exposure to pesticides affects mating, a key life history event shared by nearly all insects, and whether these impacts are mediated via impaired sexual communication. In insects, mate location and copulation are primarily regulated through chemical signals and rely on both the production and perception of semiochemicals. We show through behavioral bioassays that mating success is reduced in bumble bee gynes after exposure to field-relevant sublethal doses of imidacloprid, and that this effect is likely mediated through a disruption of both the production and perception of semiochemicals. Semiochemical production was altered in gyne and male cuticular hydrocarbons (CHCs), but not in exocrine glands where sex pheromones are presumably produced (i.e., gyne mandibular glands and male labial glands). Male responsiveness to gyne mandibular gland secretion was reduced, but not the queen responsiveness to the male labial secretion. In addition, pesticide exposure reduced queen fat body lipid stores and male sperm quality. Overall, the exposure to imidacloprid affected the fitness and CHCs of both sexes and the antennal responses of males to gynes. Together, our findings identify disruption of chemical signaling as the mechanism through which sublethal pesticide exposure reduces mating success.

## Introduction

Pesticide exposure is a major contributing factor to insect decline, including beneficial insects such as pollinators (Potts *et al*. 2010; Cameron *et al*. 2011; Goulson *et al*. 2015; Rundlöf *et al*. 2015; Brühl & Zaller 2019; Forister, Pelton & Black 2019; Sánchez-Bayo & Wyckhuys 2019). Negative impacts on pollinators are of particular concern because of their integral role in food production and wild ecosystems (Klein *et al*. 2007; Aizen *et al*. 2009; Ollerton, Winfree & Tarrant 2011; Potts *et al*. 2016; Jordan *et al*. 2021). While the effects of exposure are dependent on the dose, active ingredient, route of exposure, clearance rate (Cresswell *et al*. 2014), and species (Cresswell *et al*. 2012; Gradish *et al*. 2018; Sgolastra *et al*. 2018; Franklin & Raine 2019; Rondeau *et al*. 2022), continually mounting evidence supports the idea that even sublethal doses of pesticides negatively impact insect behavior and physiology (Blacquière *et al*. 2012; Bantz *et al*. 2018; Müller 2018; Tosi *et al*. 2022).

Globally, one of the most widely used class of pesticides are neonicotinoids (Douglas & Tooker 2015), which are applied as foliar sprays, soil drenches, and seed treatments (Jeschke *et al*. 2011) to suppress agricultural pests. After application, neonicotinoids act systemically in plants and persist in the soil, pollen and nectar, thereby creating multiple routes of exposure for pollinators (Goulson *et al*. 2015; Raine & Rundlöf 2024). Neonicotinoids disrupt central nervous system function in insects by binding to nicotinic acetylcholine receptors (Jeschke *et al*. 2011; Moffat *et al*. 2016), leading to a vast array of detrimental impacts on behavior and physiology. In bees, the negative impacts of sublethal neonicotinoid exposure include, but are not limited to: flight ability (Tosi, Burgio & Nieh 2017), foraging (Gill & Raine 2014; Colin *et al*. 2019), hygienic behavior (Tsvetkov *et al*. 2017), thermoregulation (Crall *et al*. 2018), sleep (Tasman, Rands & Hodge 2020), learning and memory (Stanley, Smith & Raine 2015; Muth, Francis & Leonard 2019; Siviter & Muth 2022), mating (Chen *et al*. 2024) and sperm viability (Chaimanee *et al*. 2016; Ciereszko *et al*. 2016; Straub *et al*. 2016). Many of these impacts likely occur simultaneously, therefore it is not surprising that sublethal neonicotinoid exposure ultimately impacts fitness by reducing reproductive output (Laycock *et al*. 2012; Whitehorn *et al*. 2012; Sandrock *et al*. 2014; Baron *et al*. 2017; Wu-Smart & Spivak 2018; Siviter *et al*. 2020; Stuligross & Williams 2020; Willis Chan & Raine 2021). However, despite the intensive efforts to describe the impacts of sublethal pesticide exposure in insects, the mechanisms mediating these effects remained poorly understood. Here, we examine how pesticide exposure affects mating, a key life-history bottleneck shared by the majority of insects, and whether the impacts are mediated through disruption of sexual communication.

Mate location and copulation in insects are primarily regulated through chemical signals called pheromones (Ayasse, Paxton & Tengö 2001; Keeling, Plettner & Slessor 2004; Yew & Chung 2015). Pheromones are often produced in exocrine glands, secreted by either males, females or both (Keeling, Plettner & Slessor 2004; Bruschini, Cervo & Turilazzi 2010) and elicit a response in the recipient. In bumble bees, males mark mating sites with secretions from their labial glands to attract gynes (newly-emerged queens) (Bergman & Bergström 1997; Ayasse & Jarau 2014). When gynes find males, short-range attraction is likely mediated by both cuticular hydrocarbons and pheromones from the head of gynes (Krieger *et al*. 2006), possibly from their mandibular glands. Previous studies suggest that this type of sexual communication can be impaired by sublethal doses of pesticide, a phenomenon known as info-disruption (Lürling & Scheffer 2007; Tricoire-Leignel *et al*. 2012). For example, in the parasitoid *Nasonia vitripennis*, imidacloprid impaired olfaction, which reduced the recognition of sexual signals and ultimately successful mating (Tappert *et al*. 2017). In a study of three tortricid moth species, sublethal exposure to thiacloprid altered female calling behavior in all three species but affected the chemical blend of the pheromone in only one species (Navarro-Roldán & Gemeno 2017). Because females of many insect species have low mating frequencies and are often singly mated (Ridley 1990; Schmid-Hempel & Schmid-Hempel 2000; Ayasse, Paxton & Tengö 2001; Bird *et al*. 2024), any reduction in their ability to find mates and successfully copulate could reduce their fitness to zero (Van Wilgenburg, Driessen & Beukeboom 2006; Heimpel & de Boer 2008). This disruption can result in large consequences on insect fitness.

Here, we examine the sublethal effects of the neonicotinoid imidacloprid on mating behavior and their underlying chemical and physiological mediators. We used *Bombus impatiens*, a native, social, and commercially available bumble bee species. *B. impatiens* is an excellent model to address these questions because like many other pollinators (mostly solitary bees), bumble bee gynes have a low mating-frequency (Schmid-Hempel & Schmid-Hempel 2000; Bird *et al*. 2024), typically mating once prior to entering a winter diapause, after which they start a social colony (Plowright & Laverty 1984). Single mating places greater importance on mate choice for gynes. Bumble bee males employ one of three mate-finding strategies: flying a patrol route between locations marked with their cephalic labial gland secretion, waiting at the nest entrance of a colony for gynes to leave, and perching and pouncing on passing gynes (Valterová *et al*. 2019). Patrolling behavior is the most common, and has been observed in many members of the *Pyrobombus* subgenus to which *B. impatiens* belongs (Svensson & Bergstrom 1977). The labial gland scent marks are attractive to gynes (Lecocq *et al*. 2015; Kubo, Harano & Ono 2017). Gynes are also assumed to produce sex pheromones in glands located in their head (Krieger *et al*. 2006), potentially the mandibular glands. These compounds presumably mediate the short-range attraction between gynes and males. We hypothesize that sublethal pesticide exposure will have detrimental effects on mating by either 1) interfering with chemical signal production or perception, and/or 2) by reducing fitness such that the insects communicate their decreased quality through honest chemical signals. To test these hypotheses, we first examine the effect of sublethal exposure on mating behavior by exposing either gynes or males to one of two sublethal doses of imidacloprid (6 ppb or 60 ppb). Next, we examine the impact of this exposure on signal production and perception through chemical and electrophysiological analyses. Finally, we measure potential fitness outcomes of imidacloprid exposure by measuring lipid storage in gynes, and sperm viability in males. Gynes rely on lipid reserves to complete diapause (Hahn & Denlinger 2007), which are accumulated in the pre-diapause period (Treanore & Amsalem 2020).

## Methods

### Bee colonies and sampling

Bees were collected from *B. impatiens* colonies obtained from Koppert Biological Systems (Howell, Michigan, USA) and housed in walk-in environmental chambers maintained at a constant temperature of 28–30°C, 60% relative humidity, and constant darkness. These colonies provided gynes (unmated, pre-diapause queens) and males, which were sampled on the day of emergence and kept separately in small plastic cages (11 cm diameter × 7 cm height). Males and gynes were grouped by colony and age cohort, to ensure they were unmated prior to experimentation. Colonies and cages were supplied with *ad libitum* 60% sucrose solution and honeybee-collected pollen. With some exceptions, the head width and wet mass of the bees used in experiments were taken as a proxy for size (Amsalem *et al*. 2009), which was used as a covariate in statistical models where applicable.

### Pesticide treatment

Imidacloprid was provided to gynes and males in all experiments listed below via chronic, *ad libitum* feeding in sugar solution. Imidacloprid (Pestanal, Sigma-Aldrich) was first diluted in deionized water to an initial concentration 100 ng/µL. This solution was further diluted in 60% sucrose solution to achieve the desired concentration. Bees were collected upon emergence to control for their age and ensure they were unmated, then were housed in small microcolony cages of 2-4 gynes and 4-6 males per cage, and access to *ad libitum* sucrose solution. On day 3, gynes were divided among treatment groups and fed either 0, 6 or 60 ppb imidacloprid for the next 3 days. Males were treated the same, except their 3 day exposure started on day 7 after emergence. Gynes were then used in mating bioassays on day 6, and males on day 10 after emergence. This feeding regime was designed to mimic chronic environmental exposure and keep the pesticide exposure consistent between gynes and males while enabling their use in mating bioassays at optimal ages of sexual maturity (i.e., males sexually mature later than gynes; (Treanore *et al*. 2021)). This served a second purpose of eliminating the variability associated with age. Six and 60 ppb concentrations represent sublethal doses that span a common range of field realistic exposure encountered in pollen and nectar, with 60 ppb representing the higher end of reported values (Schmuck *et al*. 2001; Chauzat *et al*. 2006; Krischik, Landmark & Heimpel 2007; Krupke *et al*. 2012; Stoner & Eitzer 2012; Zioga *et al*. 2020).

### Determination of imidacloprid concentration in bees

To assess the amount of imidacloprid in bees of different pesticide treatments, imidacloprid was extracted and analyzed by UPLC-MS as in (Crall *et al*. 2018). Gynes (n = 29, day 6) and males (n = 19, day 10) were homogenized in a Fastprep-bead beater with 1 mL 25% methanol, 2 metal beads, and 20 µL of 500 nM d4-imidacloprid as an internal standard. After centrifugation at 16,000 rpm for 10 min, 900 µL of supernatant was transferred to a clean 2 mL tube containing 900 µL of 25% acetic acid. Solid phase extraction columns (SPE, DSC-18, 1mL, 50 mg, Supelco) were then used to purify the samples. After conditioning the cartridges with 1 mL methanol, washing with 1 mL water, and drying, 600 µL of sample was loaded on each cartridge. The cartridges were again washed with 1 mL water, dried and eluted with 800 µL methanol into a glass vial. This was dried, resuspended in 40 µL of 50% methanol, and 5 µL injected on a Waters Acquity UPLC connected to a Waters Xevo TQ-S mass spectrometer with a BEH C18 column (1.7 µm, 2.1*100mm) at the Huck Metabolomics Core Facility. The mobile phases were A: methanol/water 2:98 v/v with 5 mM ammonium formate and 0.1% formic acid and B: methanol/water 98:2 v/v with 5 mM ammonium formate and 0.1% formic acid. The gradient started with 0% B, then increased to 20% B at 1.1 min, 90% B at 8 min, returned to 20% by 8.1 min, where it was held until 10 min. The flow rate was kept at 0.3 mL/min and the column was heated to 40°C.

### Mating bioassays

To examine the effect of sublethal imidacloprid exposure on mating, gynes and males were video recorded in paired mating bioassays. To examine how males respond to sublethally-exposed gynes, we placed one exposed and one unexposed sister gyne in a mating arena with 6 unexposed 10-day old males, and video recorded all interactions with a smartphone (iPhone 6s) or GoPro Hero 6 for 15 minutes, or until both gynes had mated (Fig 1A). The males were unrelated to the gynes but siblings with each other. This was done twice, once with one of the gynes exposed to 6 ppb imidacloprid, and once with one of the gynes exposed to 60 ppb imidacloprid. To evaluate how male pesticide exposure affects mating, we used the same setup, but inverted the exposure, with the two gynes remaining unexposed and half of the males (3/6) receiving imidacloprid (Fig 1B). The 1:3 ratio of gynes to males was maintained to increase the probability of successful mating (Treanore *et al*. 2021). Gynes and males were taken from different colonies to remove possible effects of inbreeding avoidance and nestmate recognition (Gerloff & Schmid-Hempel 2005; Whitehorn, Tinsley & Goulson 2009). Bees were paint marked with red or white on the dorsal thorax according to treatment. The color designation for treatment and control was alternated between trials. Videos were scored for the duration of male mounting behavior, and any successful mating was recorded to calculate mating frequency between groups. ‘Mounting’ was defined as the male being attached to the gynes’ abdomen in a position that precedes copulation. ‘Mating’ was defined as the stereotypical posture of the male having fallen backwards so that he was attached to the gyne only with his genitalia, with his legs twitching in the air.

**Figure 1.**
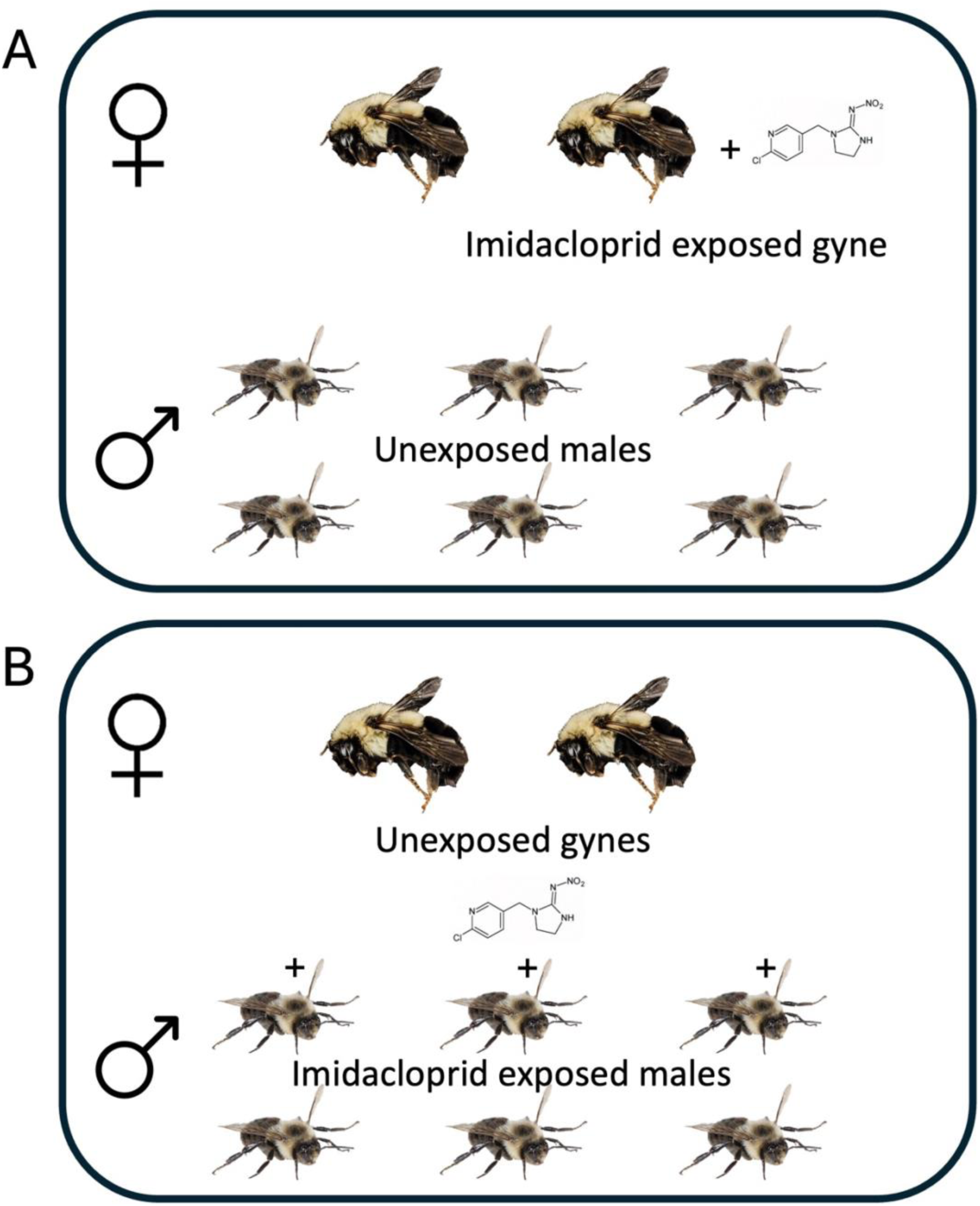
Diagram of reciprocal mating experiments. In “A” one of two gynes was fed on imidacloprid dosed sugar solution and then placed in a mating arena with six unexposed males. In “B”, three of the six males were fed imidacloprid, while the gynes were unexposed.

### Chemical extracts and analysis

All bees were frozen at -80° C prior to dissection or extraction of their cuticular hydrocarbons (CHCs) and resulting samples were stored at -20° C until analysis. Gyne mandibular glands (n = 15 per treatment) were dissected by bisecting the head with razor blade and then using a micro scissors to cut through the cuticle towards the mandible, exposing the gland. A clean, fine forceps was then used to remove the gland into a glass insert in a 2 mL glass vial containing 100 µL of dichloromethane and left to extract overnight. After removing the solvent with passive evaporation in a fume hood, 5 µL of BSTFA with 1% TCMS (Millipore Sigma) was added to make trimethylsilyl derivatives of the hydroxy acids typically found in the mandibular gland, which greatly improves their chromatography. After 4 hours at room temperature the reaction was quenched by adding 50 µL of hexane containing 1 µg of eicosane (as an internal standard) to each sample.

Male labial glands (n = 20 per treatment) were dissected by pinning the head face down in a Sylgard dissection dish and using a micro scissors to make an incision from the occipital foramen towards the vertex of the head. This incision was used as an entry point to pry up and break off pieces of cuticle and expose the labial glands, which were removed with fine forceps into a 2 mL glass vial containing 500 µL of hexane with 10 µg eicosane as an internal standard. After overnight extraction, a 100 µL aliquot was transferred to a clean glass vial with an insert and treated with BSTFA as above. 100 µL of hexane was added to quench the reaction.

CHCs of gynes and males were extracted by immersing the bees in hexane for 30 seconds. In correspondence with their body size, gynes were washed in 1.5 mL and males in 1 mL of hexane containing either 30 µg (gynes) or 20 µg (males) eicosane as internal standard.

To identify compounds, samples were analyzed on an Agilent 7890A GC equipped with a HP-5 MS column (0.25 mm id × 30 m × 0.25 µm film thickness, Agilent) and interfaced to an Agilent 5975C mass spectrometer. Compounds were tentatively identified based on retention indices and diagnostic ions, and when possible, identifications were confirmed by matching retention times and mass spectra with those of authentic standards. The double bond positions of monounsaturated cuticular hydrocarbons were determined by GC-MS analysis of their dimethyldisulfide (DMDS) adducts (Dunkelblum et al. 1985), described in Table S6 and Figure S1. DMDS derivatizations were carried out by adding 100 µl DMDS and 1 drop of 60 µg/µL I_2_ in diethyl ether to the evaporated CHC residue extracted from a single bee. The reaction was kept in a 2 mL screw-cap vial overnight at 40° C, after which 200 µl of hexane was added, and the mixture washed with an equal volume of 5% aqueous sodium thiosulfate. The organic layer was removed and dried over anhydrous sodium sulfate for GC-MS analysis.

To quantify relative and absolute amounts of compounds, 1 µL of extracts was injected on a Trace 1310 GC (Thermo Fisher, Waltham, MA USA) equipped with a flame-ionization detector (FID) and a TG-5MS column (0.25 mm id × 30 m × 0.25 µm film thickness; Thermo Fisher). The temperature program was as follows: 60°C to 120°C at 15°C/min followed by 120°C to 300°C (5 m hold) at 4°C/min. The injector port and FID were held constant at 250°C and 320°C, respectively. Analyses were conducted with Chromeleon version 7.2 SR4.

### Electroantennography

To examine the impact of imidacloprid exposure on signal perception, antennal responses to gland extracts (i.e., gyne antenna vs. male labial glands, and male antenna vs. gyne mandibular glands) were recorded using electroantennography (EAG) as previously described (Derstine *et al*. 2021), with slight modifications. The antennae of gynes (total sample size = 60; control - 30, 6 ppb – 15, 60 ppb - 15) and males (total sample size = 54; n = 18 per treatment group) fed with 0, 6, and 60 ppb imidacloprid were exposed to a control stimulus and a gland extract from the opposite sex. Mandibular glands from 70 gynes and labial glands from 50 males were dissected into hexane to create pooled extracts that ensured consistent stimuli across replicates. The pooled extracts were evaporated to a concentration of 1 gland equivalent per 10 µL. Odor cartridges were created by pipetting 1-bee equivalent of either mandibular or labial extract, or 10 µL hexane as a blank control, onto strips of Whatman filter paper. These strips were inserted into glass pipettes after the solvent evaporated and connected to the air delivery tube. For EAG preparations, a single antenna was excised, and the tip snipped with fine scissors prior to mounting between the paddles of a Syntech EAG probe with Spectra 360 electroconductive gel and placed in the air stream. Each antenna was allowed to reach a steady baseline before starting the experiment, at which point the test odorants were delivered into a steady airstream by using a stimulus flow controller (Syntech, Hilversum, Netherlands) to pass air through the odor cartridges. Responses to each stimulus were amplified with a 10 × pre-amplifier (Syntech, Netherlands) and further amplified 10 × before being recorded and analyzed on a desktop computer using Syntech EAG software.

Each antenna was exposed to two control, and two gland extract puffs, all 15 seconds apart. For each antenna, responses to test odorants were averaged and normalized by subtracting the average response to the control odorants and the corrected values were used in analyses.

### Lipid measurement

Total lipids per mass of gynes’ fat body was measured with the vanillin-phosphoric acid bioassay as in (Treanore & Amsalem 2020). Eviscerated abdomens (control n = 30, 6 ppb n = 27, 60 ppb n = 32) were weighed and then homogenized in 1 mL 2% sodium sulfated solution. For each sample, a 200 µL aliquot of the homogenate was added to 2.8 mL of 1:1 chloroform:methanol (v:v), vortexed for 30 seconds, and centrifuged to precipitate glycogen (3000 rpm for 5 min). The supernatant was transferred to a clean glass vial and mixed with an additional 2 mL distilled water. The upper fraction was discarded while the lower fraction containing lipids was reserved. After removing the solvent by evaporation, 200 µL sulfuric acid was added to the lipid residue and heated for 10 min at 100°C. 5 mL of vanillin-phosphoric acid reagent was added, and after cooling, 300 µL was transferred to a 96-well plate in triplicate. Absorbance was measured at 525 nm using a BioTek Microplate Spectrophotometer (BioTek Instruments Inc., Winooski, VT). For each sample, resulting absorbance values measured against a standard curve which consisted of five different concentrations of 0.1% vegetable oil diluted in chloroform. Absorbance values were converted to micrograms based on a formula calculated from the regression line derived from the standard curve. Lipid quantity is presented as lipids in µg per mg of fat body to correct for body size.

### Sperm viability

Sperm from 10-day old males exposed to 0 ppb, 6 ppb, and 60 ppb imidacloprid (n = 17 per treatment group) were visualized using an Olympus Fluoview 10i-LIV confocal microscope after staining with the Live/Dead Sperm Viability Kit (L-7011). This kit uses SYBR 14 and propidium iodide to fluorescently stain live and dead sperm, respectively. The dyes were diluted according to manufacturer’s instructions in Kiev buffer and used on the day of preparation. Males treated with 0, 6, or 60 ppb imidacloprid dosed sucrose solution were dissected at 10 days old after cold anesthesia. Accessory testes were removed into a 25 µL drop of Kiev buffer, and gently torn open with forceps to allow sperm to flow out into the liquid. The liquid was transferred to a 2 mL Eppendorf tube containing 25 µL Live/Dead dye working solution and incubated at 27° C for 10 minutes. 5 µL of stained sperm solution was pipetted onto a clean glass slide, covered with a glass coverslip, and scanned in green and red wavelengths. From the resulting image, 10 subsections were randomly selected, and all live and dead sperm were counted within each subsection to calculate the proportion of live sperm. These subsections were also averaged to give the mean total sperm per sampling area.

### Statistics

Statistical analyses were performed using R (version 4.1.3) in RStudio. In mating trials, male mounting behavior was analyzed with a paired Wilcoxon Rank Sum test, whereas the proportion of successful mating between control and treatment groups was analyzed using a two-proportion Z-test. Permutational analysis of variance (PERMANOVA) was used to examine overall chemical differences in extracts of gyne mandibular glands, male labial glands, and CHCs between treatment groups of gynes and males. Only peaks > 0.5% of the total peak area were used. These selected peaks were then re-standardized to 100%. Individual components were compared using linear mixed effects models with colony as a random factor and Tukey’s Honest Significant Difference post-hoc test to obtain pairwise contrasts of treatment groups. In the analyses of EAG, total lipids, and sperm viability, we used linear mixed effects models with colony as a random effect to test the impact of treatment on the response variable. Pairwise contrasts between treatment groups were subsequently evaluated with Tukey’s Honest Significant Difference post-hoc test.

## Results

### Imidacloprid body concentration

Gynes and males consuming sucrose solution containing imidacloprid for 3 days had comparable amounts of imidacloprid in their body when given 6 ppb (gynes - 0.093 ng ± 0.02, males - 0.091 ng ± 0.01, mean ± SE, n = 10) and 60 ppb (gynes - 1.005 ng ± 0.31, males - 1.102 ng ± 0.18, mean ± SE, n = 10), while imidacloprid was not found in any of the control gynes (Fig 2). The statistics of all pairwise comparisons are given in Table S1.

**Figure 2.**
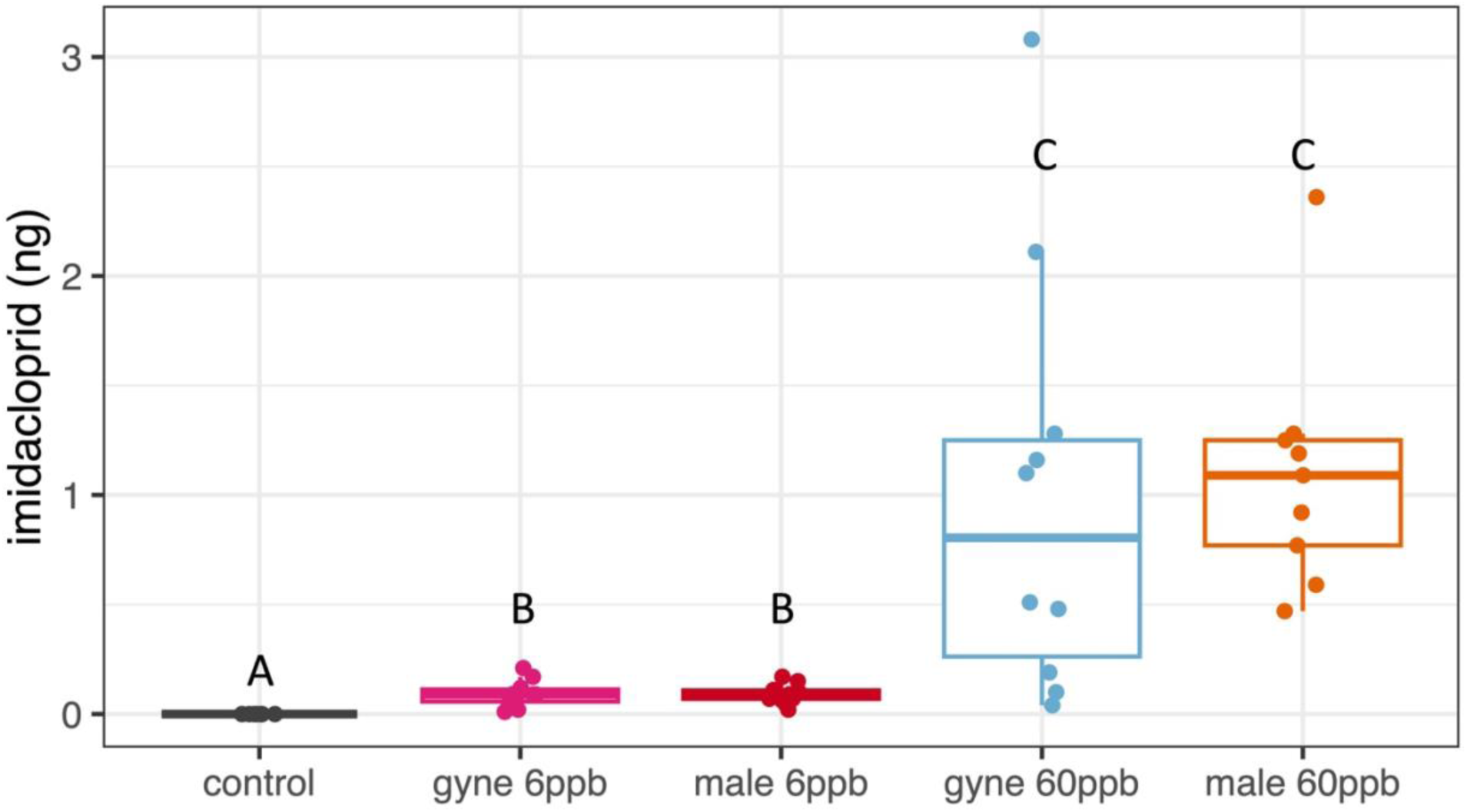
LCMS analysis of imidacloprid amount in individual gynes and males after 3 days of feeding on either control, 6 ppb, or 60 ppb imidacloprid-dosed sugar solution. Letters above the boxplots indicate statistical differences determined by pairwise contrasts using the Kruskal-Wallis test followed by Dunn’s post hoc test with Benjamini and Hochberg adjusted p-values (α = 0.05).

### Effect of imidacloprid on mating behavior

Treatment with 6 ppb imidacloprid reduced mating attempts by males toward exposed gynes (Fig 3). Gynes which consumed 6 ppb imidacloprid received less male mounting (p = 0.008, Wilcoxon Rank Sum test; 1.2 ± 0.6 and 3 ± 0.7 minutes, mean ± SE in 6 ppb vs. control groups, respectively, Fig 3A) and mated less frequently than unexposed gynes (p = 0.05 two-proportion Z-test, n = 30, 20% to 43% in 6 ppb vs. control groups, respectively, Fig 3B). In the 60 ppb treatment group, the overall mating success was significantly lower in both the treatment and control groups and there were no significant differences between treatment groups in male mounting (p = 0.97, Wilcoxon Rank Sum test, 0.27 ± 0.2 min and 0.45 ± 0.2 min in 6 ppb vs control groups, respectively, Fig 3C) or mating frequency (p = 0.09 two-proportion Z-test, n = 30, 3% to 14% in 6 ppb vs. control groups, respectively, Fig 3D).

**Figure 3.**
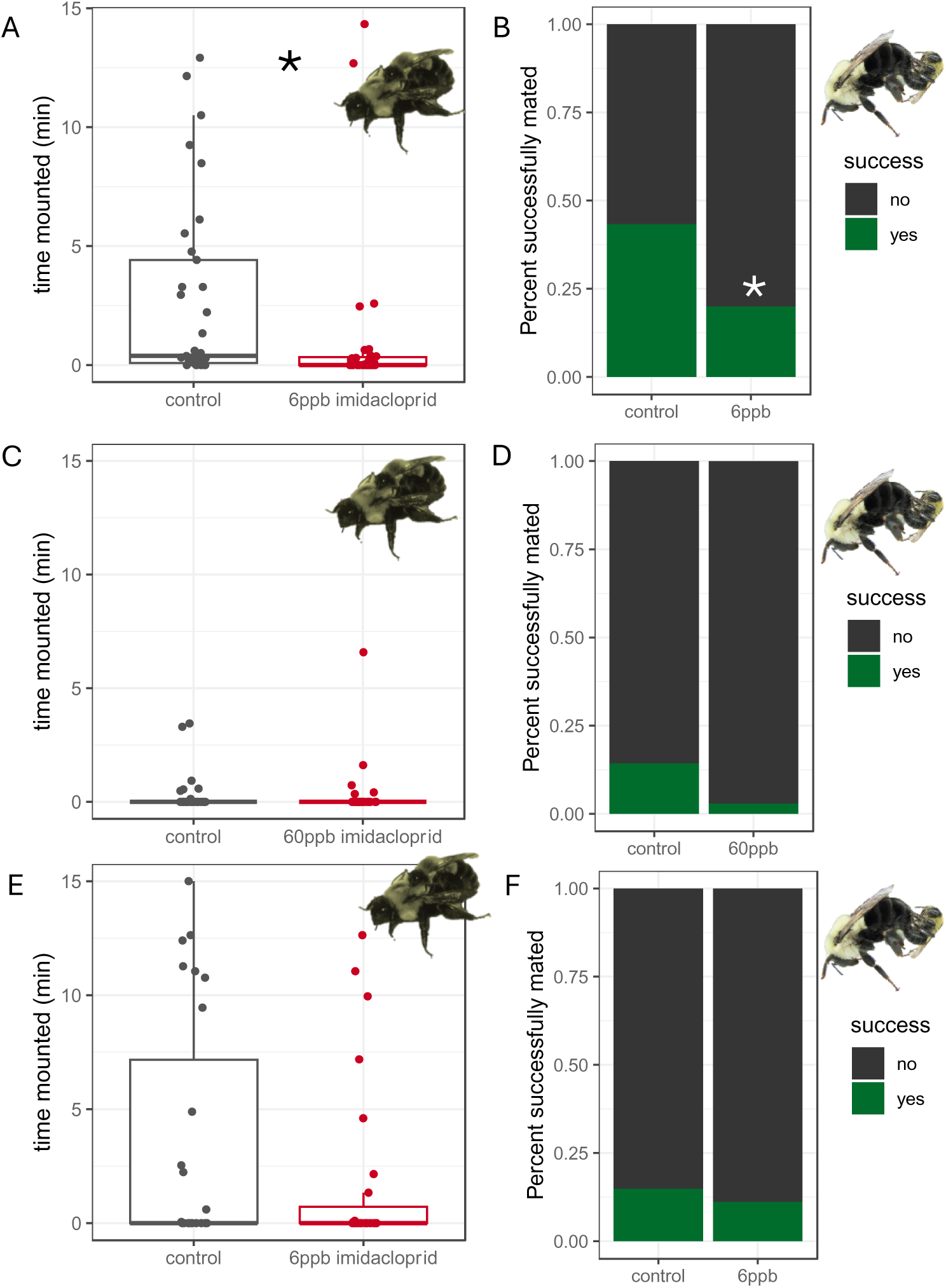
The impact of imidacloprid exposure on mounting behavior by males (A,C,E) and mating behavior (B,D,F) when either gynes or males were fed imidacloprid. Panels A-D correspond to mating trials where half of the gynes were fed imidacloprid and, panels E-F correspond to mating trials where half of the males were fed imidacloprid. Asterisks denote significant differences at p < 0.05.

In mating trials where only the males were exposed to imidacloprid, there was no effect of treatment on male mounting (p = 0.19, Wilcoxon Rank Sum test; 1.8 ± 0.7 and 3.4 ± 1.0 minutes, mean ± SE in 6 ppb vs. control groups, respectively, Fig 3E) or successful mating (p = 0.68 two-proportion Z-test, n = 27, 11% to 14% in 6 ppb vs control groups, respectively, Fig 3F).

### Effect of imidacloprid on chemical signaling

To evaluate if altered chemical signaling was responsible for reduced mating success in bees exposed to imidacloprid, we examined CHCs in gynes and males, gyne mandibular glands and male labial glands in both control and treatments. Gyne and male CHCs were comprised primarily of saturated and unsaturated aliphatic alkanes and alkenes, with gynes having a much higher proportion of unsaturated compounds than males. These desaturations occurred predominately at the 9 and 7 positions for all chain lengths encountered. Gyne mandibular glands contained a series of 3-hydroxy acids in addition to hydrocarbons and several compounds which did not vary across treatments and thus their identification was not pursued. Male labial glands contained predominantly two compounds, 2,3-dihydrofarnesol (previously reported in (Luxová *et al*. 2004)), and hexadecenoic acid, in addition to a smaller proportion of hydrocarbons and farnesyl esters. As in the mandibular glands, because these compounds were not affected by the pesticide treatment, their complete identification was not pursued. The amounts of unidentified compounds were included in all analyses, provided they met the 0.5% relative peak area cutoff. The identity, retention indices, and average proportion of compounds used in statistical analyses are given in Table S2 (Gyne CHCs), Table S3 (Male CHCs), Table S4 (Gyne mandibular gland), and Table S5 (Male labial gland).

Gyne CHCs were altered by 60 ppb but not 6 ppb imidacloprid exposure. An NMDS plot of gyne CHCs (Fig 4A) shows separation between the 60 ppb imidacloprid group and the two other groups (0 and 6 ppb) which cluster together. This visualization is supported by pairwise PERMANOVA which finds CHCs in the 60 ppb group significantly different from the 6 ppb (p = 0.006) and the control (p = 0.05) groups. One compound in particular seems to drive this separation, 9-pentacosene (Fig 4B). Simultaneous pairwise comparisons using a linear mixed effects model with Tukey’s HSD test indicated 9-pentacosene was significantly increased in the 60 ppb group, representing 18.1 ± 0.5% of the total peak area in the 60 ppb treatment group, compared with 15.6 ± 0.6% in the 6 ppb treatment group and 15.3 ± 0.4% in the control group (mean ± SE, n = 15; 60 ppb vs. 6 ppb p = 0.001, 60 ppb vs. control p = 0.0001, 6 ppb vs. control p = 0.88).

**Figure 4.**
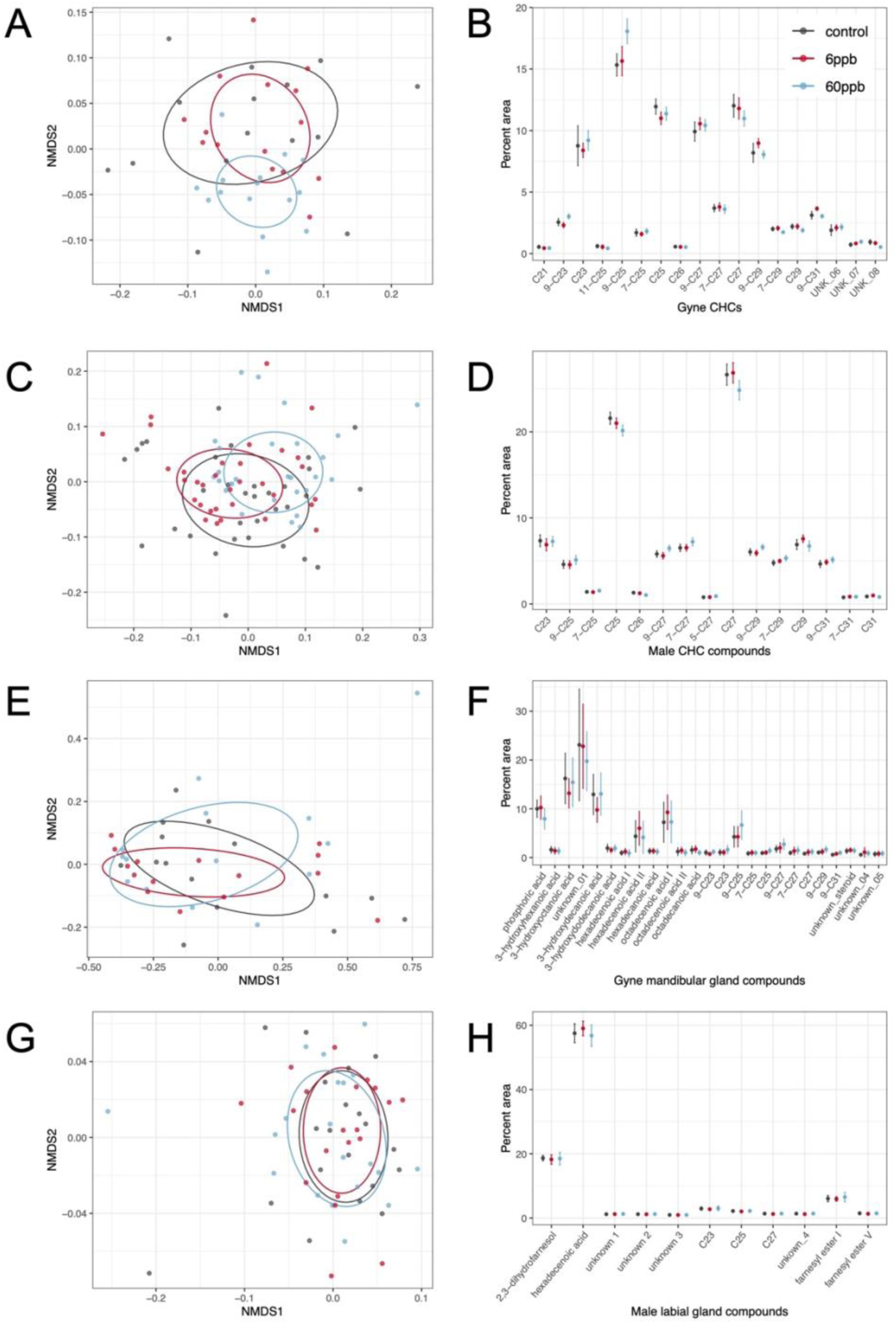
The effect of pesticide exposure on signal production. Panels A, C, E, G show NMDS plots based on the Bray-Curtis dissimilarity matrix of relative amounts of chemical components in gland extracts or cuticular washes. Panels B, D, F, H show relative amounts of each chemical component quantified from GC-FID and GC-MS analyses. Points are means ± 2SE (to approximate 95% confidence intervals). Panels A,B and C,D describe gyne and male cuticular hydrocarbon data, respectively. Panels E,F and G,H describe gyne mandibular gland and male labial gland extracts, respectively.

Similarly, exposure to 60 ppb, but not 6 ppb imidacloprid, altered the overall male CHC profile (Fig 4C, pairwise PERMANOVA, 60 ppb vs. 6 ppb p = 0.01, 60 ppb vs. control p = 0.02, 6 ppb vs. control p = 0.36). However, this difference was not obviously explained by individual compounds, and while the entire profile differed between the groups, no single individual compound was significantly different between treatment groups (Fig 4D).

In contrast, imidacloprid exposure had no effect on the chemical composition of the gyne mandibular gland (p = 0.81, PERMANOVA, Fig 4E-F) or the male labial gland (p = 0.45, PERMANOVA, Fig 4G-H), at either 6 ppb or 60 ppb doses.

### Effect of imidacloprid on signal perception

To evaluate if altered perception of chemical signaling was responsible for reduced mating success in bees exposed to imidacloprid, we examined gyne perception of male labial gland secretion and male perception of gyne mandibular gland secretion. EAG recordings of gynes responding to male labial gland extract showed that imidacloprid did not affect signal perception at the tested doses (Fig. 5A, B). A linear mixed effects model found no significant differences of 6 ppb treatment (χ^2^ = 0.56, *df* = 1, p = 0.45), or 60 ppb treatment (χ^2^ = 0.004, *df* = 1, p = 0.98) compared to their corresponding controls. However, signal perception of males responding to gyne mandibular gland extract was affected by the imidacloprid treatment (Fig 5C, χ^2^ = 7.7, *df* = 2, p = 0.02). Post-hoc comparisons between treatment groups showed that males in the 60 ppb, but not in the 6 ppb group, had significantly lower EAG responses than controls (Tukey HSD, 60 ppb vs. control p = 0.02, 60 ppb vs. 6 ppb p = 0.11, 6 ppb vs. control p = 0.93).

**Figure 5.**
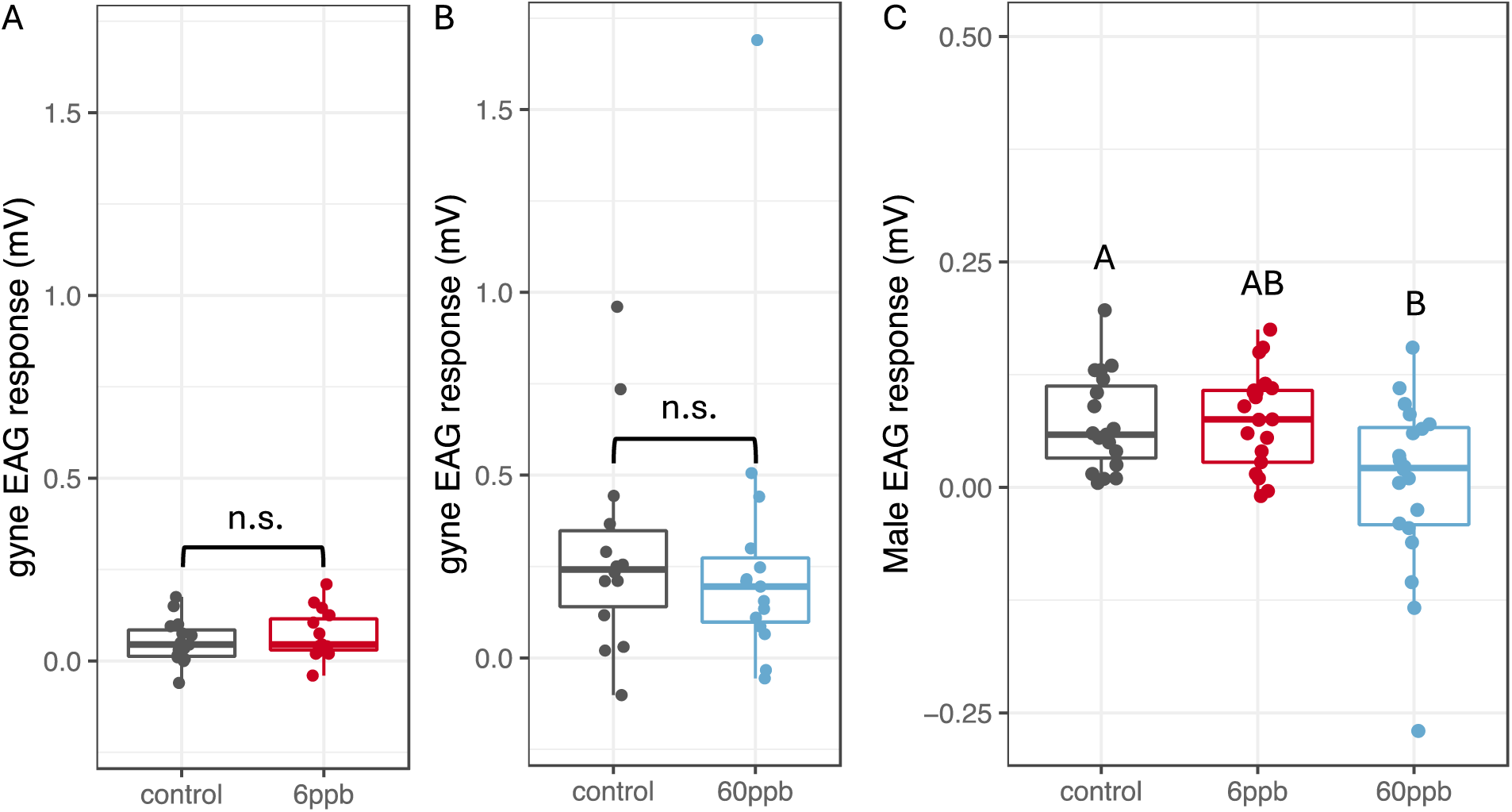
The effect of imidacloprid exposure on signal perception. Panels show antennal responses of gynes (A, B) and males (C) exposed to either control, 6 ppb, or 60 ppb imidacloprid after receiving puffs of male labial gland extract. As the 6 ppb and 60 ppb treatments were run separately for gynes, they are presented separately and next to their concurrently run controls. Letters above boxplots denote statistical differences of pairwise contrasts at p < 0.05 (calculated from a linear mixed effects model with colony as a random factor, using Tukey’s HSD post-hoc test for pairwise comparisons).

### Effect of imidacloprid on gyne fat body lipid stores

Imidacloprid exposure at the 60 ppb, but not 6 ppb dose, reduced gyne fat body lipid stores compared to controls (Fig 6, χ^2^ = 15.5, *df* = 2, p < 0.001; 133.4 ± 8.2, 115.1 ± 7.4 and 95.5 ± 4.3 µg lipid/mg fat body ± SE, n = 30/27/32 for 0, 6 and 60 ppb, respectively). Post-hoc comparisons between treatment groups showed that the 60 ppb, but not the 6 ppb group, had significantly lower fat body lipids than controls (Tukey HSD, 60 ppb vs. control p < 0.001, 60 ppb vs. 6 ppb p = 0.19, 6 ppb vs. control p = 0.09).

**Figure 6.**
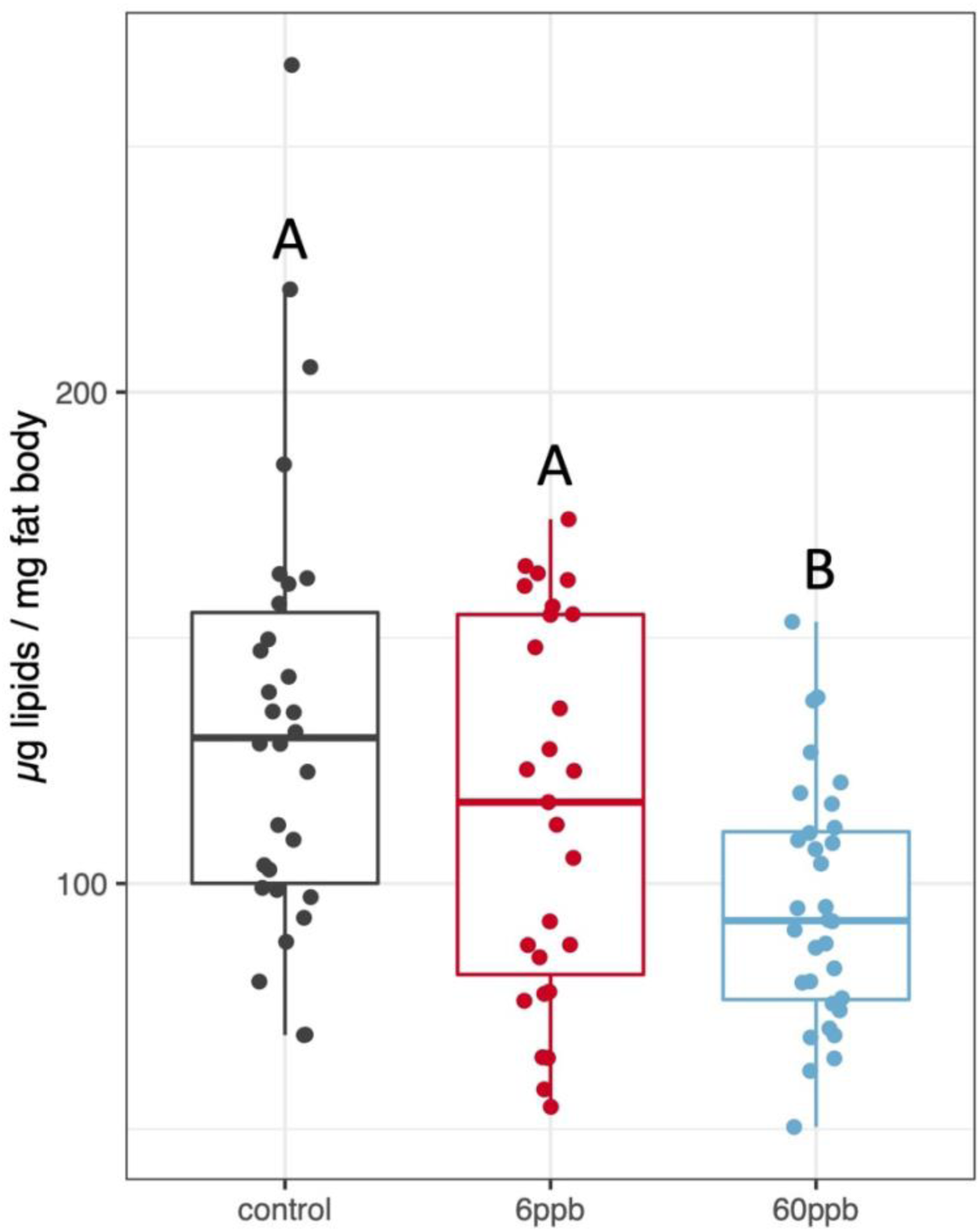
Lipid stores of 6-days old gynes fed either 0, 6, or 60 ppb imidacloprid in sugar solution. Values are presented in micrograms of lipid per milligram of fat body tissue. Letters above boxplots indicate significant differences (linear mixed effects model followed by Tukey HSD, α = 0.05).

### The effect of imidacloprid on male sperm viability

Imidacloprid exposure in males affected sperm viability in the 60 ppb but not in the 6 ppb group (Fig 7). The mean total number of sperm was different between groups (χ^2^ = 9.4, *df* = 2, p = 0.01), of which the 60 ppb group was reduced compared to controls (Tukey HSD: 60 ppb vs. 6 ppb p = 0.01, 60 ppb vs. 6 ppb p = 0.07, 6 ppb vs. control p = 0.71). The mean proportion of live sperm was not significantly different between treatment groups (χ^2^ = 5.4, *df* = 2, p = 0.07; Fig 7B).

**Figure 7.**
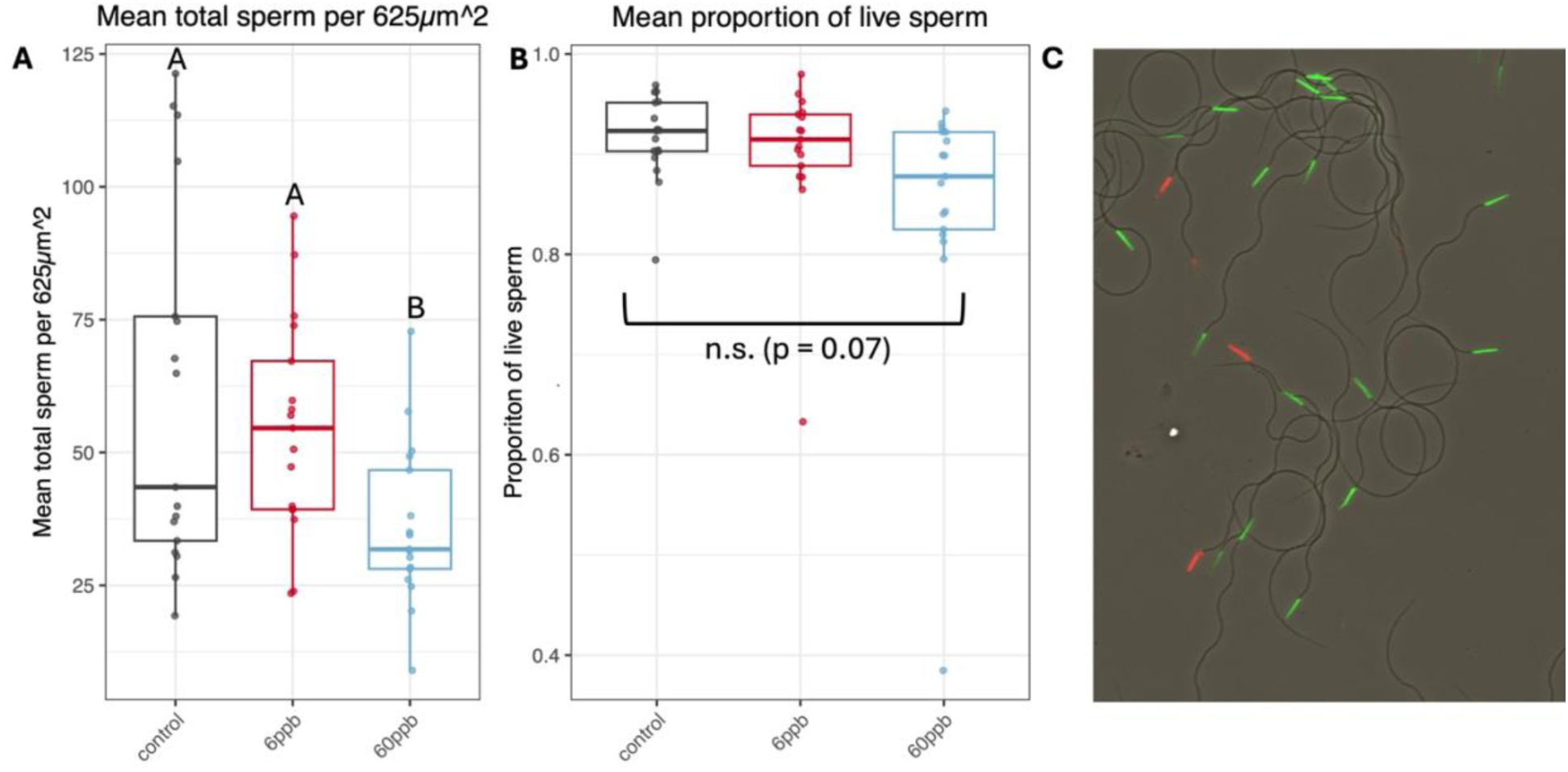
A – The mean number of sperm counted per subsection, per male. B – The proportion of live sperm in each treatment group. Letters above boxplots indicate significant differences (linear mixed effects model followed by Tukey HSD, α = 0.05). C – Example image of fluorescently dyed *B. impatiens* sperm (green indicates living and red indicates dead sperm).

## Discussion

Our results show that chronic sublethal exposure to imidacloprid negatively affects mating behavior in *B. impatiens* at doses as low as 6 ppb and identifies altered semiochemical production and perception as a likely mechanism mediating this effect. The effect on mating was sex specific and the gynes were more vulnerable to the same dose compared to the males, despite being 2-3 times larger than the males. Imidacloprid exposure altered the cuticular hydrocarbon profile of both sexes and reduced male antennal sensitivity to the gyne mandibular glands at the 60 ppb dose. Furthermore, we found negative effects on sperm viability in males and lipid storage in gynes when exposed to 60 ppb imidacloprid. Given that mate location and copulation are primarily regulated by chemical signaling in insects, the effects of pesticides on chemical communication adds to growing concerns about the impact of widespread pesticide use on insect populations (Forister, Pelton & Black 2019; Sánchez-Bayo & Wyckhuys 2019; Main *et al*. 2020; Minnameyer *et al*. 2021; Willis Chan & Raine 2021).

While the 60 ppb dose negatively affected nearly all the chemical and physiological variables we tested, including chemical production and perception, total lipids, and sperm viability, it did not influence mating behavior. This outcome contrasts with the negative impact on mating behavior observed at the 6 ppb dose. We suspect that the lack of behavioral impact at the 60 ppb dose is an artifact of the overall lower mating frequency in that experiment, which likely reduced statistical power and increased the risk of false negative (Type II error). Overall, bees clearly detected and responded to changes in the 6 ppb treatment that were below the sensitivity of our instruments but became evident with the higher 60 ppb dose.

Our data further indicate a stronger effect of the same dose on gynes compared to males. This is evident in two key findings: first, gynes had comparable pesticide levels in their bodies to males, despite being 2–3 times larger; and second, the treatment had an asymmetrical impact on mating behavior, with males avoiding treated gynes, while gynes showed no such avoidance of treated males. Sex differences in response to pesticide exposure have several possible explanations. Males and females of many insect species have clear differences in overall body size and the amount of fat body tissue, and in bees, also in the number of chromosomes. All of these could interact to affect the processing and physiological response to pesticide exposure. In *B. impatiens,* gynes can weigh 2-5x more than males (see Fig S2), which could impact both the ultimate body concentration of pesticide and how much nectar is consumed. Once ingested, imidacloprid is primarily metabolized rather than excreted (Suchail *et al*. 2004; Suchail, Debrauwer & Belzunces 2004). This is significant because detoxification enzymes are predominantly produced in fat body tissue (Li, Yu & Feng 2019), which is much more abundant in gynes than males (yet, was not more effective in gynes vs. males in our data). Additionally, gynes are diploid. Haploid individuals can be more susceptible to neonicotinoids than their diploid sisters, likely because of reduced variation in detoxification genes (Friedli *et al*. 2020). The combined effects of these interactions argue that males should suffer greater effects from pesticide exposure than females. However, in our experiments and others (Boff & Ayasse 2023), sublethal pesticide exposure did not significantly reduce mating when only males were exposed, as it did when only females were exposed. In addition, the concentration of pesticides provided and the total amount found in the body of males and gynes was the same, despite differences in body size. Thus, it is more likely that the differences in behavior of pesticide exposed males and gynes found in this study are a result of differences in the level of choosiness by gynes and males and the range at which mate choice takes place. This is in line with the assumption that bumble bee gynes choose males from a long range by attracting to their labial gland secretion and mate choice in the short range is likely determined by the males. Differences in pesticide levels in the body could be influenced not only by body size but also by consumption, as the pesticides were administered chronically. However, sugar consumption was not measured in this study, so we cannot determine whether males and gynes arrived at similar amounts of imidacloprid in the body because they regulated their consumption differentially, or if they had different clearance rates. This would be a worthwhile subject of future investigation.

Given the mating system of *B. impatiens*, where males can mate multiply but gynes rarely do (Starr 1984), gynes should be more incentivized to choose high quality mates than males (Boomsma, Kronauer & Pedersen 2009). In this study, gynes were less selective than males. Generally, it is assumed that long distance mate-finding in *Pyrobombus* bumble bees is mediated by female attraction to male labial gland pheromones (Valterová *et al*. 2019), and that less volatile CHCs may play a role once contact is initiated. Thus, females have an opportunity to assess and choose males both in the initial long-range attraction phase, and in close pre-copulatory behavior, while male choice can only be exercised in the later. One possible explanation for the data presented here is that by placing males and gynes together in a mating arena, the long-distance mate finding step mediated by male labial gland pheromones was circumvented, preventing one avenue of mate discrimination or assessment (by gynes), which translates into negative impacts on mating in that sex.

Under honest signaling theory, chemical signals are expected to convey true and reliable information in order to be evolutionarily stable (Smith & Harper 2003). In moths, for example, sex pheromone signals have been shown to covary with fitness and convey information about individual quality (Foster & Johnson 2011; Blankers *et al*. 2021). In this context, pheromones should convey accurate information about mate quality. However, the putative volatile sex pheromone compounds of both sexes were unaffected by imidacloprid treatment at doses which impacted clear components of what would make a quality mate - sperm viability in males and lipid stores in gynes. This would suggest that these signals are not honest, or at least are not tied to these measures of quality and warrants more research to identify the specific compounds responsible for sex attraction in *B. impatiens.* Instead, the profile of cuticular hydrocarbons of both males and females was altered when bees ingested imidacloprid, potentially acting as honest signals.

It is important to note that while the male labial glands were shown to contain a sex pheromone across multiple species of bumble bees (Valterová *et al*. 2019), the production of a sex pheromone in the gyne mandibular gland is not a well-established phenomenon. One study relied on a highly invasive gland excision surgery with low replication and variable results in different species (van Honk, Velthuis & Röseler 1978). Another demonstrated that while a freshly frozen gyne could elicit copulation from males, extracts from both the cuticular surface and whole head (but not specifically the mandibular glands) were successful in eliciting mounting, but not copulation, from males (Krieger *et al*. 2006). These extracts contained GC-EAD active compounds but were not localized to a particular gland. Urbanová et al subsequently found many of the electrophysiologically active compounds, including the 3-hydroxy acids, in the mandibular glands (Urbanová *et al*. 2008). This indicates that the male response may require multi-modal signals to copulate, and that a whole bee extract is more likely to contain the sex pheromone, given that compounds from multiple glands may be spread onto the cuticle to elicit copulation.

Pesticides’ effects on mating and sexual communication are often elusive and hard to detect since they manifest in species- and sex-specific ways and do not always affect fitness immediately. For example, in the mason bee *Osmia cornuta,* fungicide exposure alters male vibrational and cuticular signals which decreases mating (Boff *et al*. 2022), while in another megachilid, *Heriades truncorum,* pesticide exposure affects precopulatory behaviors in males and cuticular signals females (Boff & Ayasse 2023). Similar to our findings, pesticide exposure did not affect the proportion of successful mating in *H. truncorum* in males but did in females.

Overall, our data show reduced mating success in gynes exposed to imidacloprid accompanied by changes in their cuticular profiles and the responsiveness of males to their chemical secretion. We further demonstrate fitness consequences to imidacloprid exposure in both gynes and males and together, propose a plausible mechanism explaining the impact of imidacloprid on mating with disrupt sexual communication. Such disruption to communication systems is not likely to be immediately apparent in insects and its consequences may be extended over several generations. These data add up to the growing concerns of the use of sublethal, field relevant, doses of neonicotinoids and their impacts on pollinator health. Finally, to control as many variables as possible, we chose an exposure scenario of 3 days of adult exposure, which is conservative with regards to exposure scenarios of wild bees. In a natural setting, bees might contend with pesticide exposure during development (Kozii *et al*. 2021) and potentially continuously as adults, both of which have been shown to have detrimental effects on queens.

## Acknowledgements

The authors would like to acknowledge the Huck Metabolomics Core Facility (RRID:SCR_023864) for use of the Waters Acquity UPLC and Xevo TQ-S mass spectrometer and Dr. Ashley Shay for collecting the data shown in Figure 2. We also acknowledge the Huck Microscopy Core Facility (RRID:SCR_024457) for use of the Olympus Fluoview 10i-LIV confocal microscope. This research was funded through a USDA-NIFA Predoctoral Fellowship (2021-67034-35047) to ND.

## Author Contributions

N.D. designed and carried out the experiments, analyzed the data and prepared the figures, and wrote the main manuscript text. E.A. designed experiments, wrote and revised the manuscript. Under the supervision of N.D., C.M. and F.P collected callow bees, carried out mating experiments, administered pesticide treatments, and took size measurements of bees (weight and head width).

## References

Aizen, M.A., Garibaldi, L.A., Cunningham, S.A. & Klein, A.M. (2009) How much does agriculture depend on pollinators? Lessons from long-term trends in crop production. Annals of Botany, 103, 1579–1588.

Amsalem, E., Twele, R., Francke, W. & Hefetz, A. (2009) Reproductive competition in the bumble-bee *Bombus terrestris*: do workers advertise sterility? Proc Biol Sci, 276, 1295–1304.

Ayasse, M. & Jarau, S. (2014) Chemical ecology of bumble bees. Annu Rev Entomol, 59, 299–319.

Ayasse, M., Paxton, R.J. & Tengö, J. (2001) Mating behavior and chemical communication in the order Hymenoptera. Annual review of entomology, 46, 31–78.

Bantz, A., Camon, J., Froger, J.-A., Goven, D. & Raymond, V. (2018) Exposure to sublethal doses of insecticide and their effects on insects at cellular and physiological levels. Current Opinion in Insect Science, 30, 73–78.

Baron, G.L., Jansen, V.A.A., Brown, M.J.F. & Raine, N.E. (2017) Pesticide reduces bumblebee colony initiation and increases probability of population extinction. Nat Ecol Evol, 1, 1308–1316.

Bergman, P. & Bergström, G. (1997) Scent marking, scent origin, and species specificity in male premating behavior of two Scandinavian bumblebees. Journal of Chemical Ecology, 23, 1235–1251.

Bird, S.A., Pope, N.S., McGrady, C.M., Fleischer, S.J. & López-Uribe, M.M. (2024) Mating frequency estimation and its importance for colony abundance analyses in eusocial pollinators: a case study of *Bombus impatiens* (Hymenoptera: Apidae). Journal of economic entomology.

Blacquière, T., Smagghe, G., van Gestel, C.A.M. & Mommaerts, V. (2012) Neonicotinoids in bees: a review on concentrations, side-effects and risk assessment. Ecotoxicology, 21, 973–992.

Blankers, T., Lievers, R., Plata, C., van Wijk, M., van Veldhuizen, D. & Groot, A.T. (2021) Sex pheromone signal and stability covary with fitness. Royal Society open science, 8, 210180.

Boff, S. & Ayasse, M. (2023) Exposure to sublethal concentration of flupyradifurone alters sexual behavior and cuticular hydrocarbon profile in *Heriades truncorum*, an oligolectic solitary bee. Insect Science.

Boff, S., Conrad, T., Raizer, J., Wehrhahn, M., Bayer, M., Friedel, A., Theodorou, P., Schmitt, T. & Lupi, D. (2022) Low toxicity crop fungicide (fenbuconazole) impacts reproductive male quality signals leading to a reduction of mating success in a wild solitary bee. Journal of Applied Ecology, 59, 1596–1607.

Boomsma, J.J., Kronauer, D.J. & Pedersen, J. (2009) The evolution of social insect mating systems. Organization of insect societies, 3–25.

Brühl, C.A. & Zaller, J.G. (2019) Biodiversity decline as a consequence of an inappropriate environmental risk assessment of pesticides. Frontiers in Environmental Science, 177.

Bruschini, C., Cervo, R. & Turilazzi, S. (2010) Pheromones in Social Wasps. Pheromones, pp. 447–492.

Cameron, S.A., Lozier, J.D., Strange, J.P., Koch, J.B., Cordes, N., Solter, L.F. & Griswold, T.L. (2011) Patterns of widespread decline in North American bumble bees. Proc Natl Acad Sci U S A, 108, 662–667.

Chaimanee, V., Evans, J.D., Chen, Y., Jackson, C. & Pettis, J.S. (2016) Sperm viability and gene expression in honey bee queens (*Apis mellifera*) following exposure to the neonicotinoid insecticide imidacloprid and the organophosphate acaricide coumaphos. J Insect Physiol, 89, 1–8.

Chauzat, M.-P., Faucon, J.-P., Martel, A.-C., Lachaize, J., Cougoule, N. & Aubert, M. (2006) A survey of pesticide residues in pollen loads collected by honey bees in France. Journal of economic entomology, 99, 253–262.

Chen, X., Wang, Y., Zhou, Y., Wang, F., Wang, J., Yao, X., Imran, M. & Luo, S. (2024) Imidacloprid reduces the mating success of males in bumblebees. Science of The Total Environment, 928, 172525.

Ciereszko, A., Wilde, J., Dietrich, G.J., Siuda, M., Bąk, B., Judycka, S. & Karol, H. (2016) Sperm parameters of honeybee drones exposed to imidacloprid. Apidologie, 48, 211–222.

Colin, T., Meikle, W.G., Wu, X. & Barron, A.B. (2019) Traces of a Neonicotinoid Induce Precocious Foraging and Reduce Foraging Performance in Honey Bees. Environmental science & technology, 53, 8252–8261.

Crall, J.D., Switzer, C.M., Oppenheimer, R.L., Ford Versypt, A.N., Dey, B., Brown, A., Eyster, M., Guerin, C., Pierce, N.E., Combes, S.A. & de Bivort, B.L. (2018) Neonicotinoid exposure disrupts bumblebee nest behavior, social networks, and thermoregulation. Science, 362, 683–686.

Cresswell, J.E., Page, C.J., Uygun, M.B., Holmbergh, M., Li, Y., Wheeler, J.G., Laycock, I., Pook, C.J., de Ibarra, N.H., Smirnoff, N. & Tyler, C.R. (2012) Differential sensitivity of honey bees and bumble bees to a dietary insecticide (imidacloprid). Zoology, 115, 365–371.

Cresswell, J.E., Robert, F.X., Florance, H. & Smirnoff, N. (2014) Clearance of ingested neonicotinoid pesticide (imidacloprid) in honey bees (*Apis mellifera*) and bumblebees (*Bombus terrestris*). Pest Manag Sci, 70, 332–337.

Derstine, N., Villar, G., Orlova, M., Hefetz, A., Millar, J. & Amsalem, E. (2021) Dufour’s gland analysis reveals caste and physiology specific signals in *Bombus impatiens*. Sci Rep, 11, 2821.

Douglas, M.R. & Tooker, J.F. (2015) Large-Scale Deployment of Seed Treatments Has Driven Rapid Increase in Use of Neonicotinoid Insecticides and Preemptive Pest Management in U.S. Field Crops. Environmental science & technology, 49, 5088–5097.

Forister, M.L., Pelton, E.M. & Black, S.H. (2019) Declines in insect abundance and diversity: We know enough to act now. Conservation Science and Practice, 1.

Foster, S.P. & Johnson, C.P. (2011) Signal Honesty through Differential Quantity in the Female-Produced Sex Pheromone of the Moth *Heliothis virescens*. Journal of Chemical Ecology, 37, 717–723.

Franklin, E.L. & Raine, N.E. (2019) Moving beyond honeybee-centric pesticide risk assessments to protect all pollinators. Nature Ecology & Evolution, 3, 1373–1375.

Friedli, A., Williams, G.R., Bruckner, S., Neumann, P. & Straub, L. (2020) The weakest link: Haploid honey bees are more susceptible to neonicotinoid insecticides. Chemosphere, 242, 125145.

Gerloff, C.U. & Schmid-Hempel, P. (2005) Inbreeding depression and family variation in a social insect, *Bombus terrestris* (Hymenoptera: Apidae). Oikos, 111, 67–80.

Gill, R.J. & Raine, N.E. (2014) Chronic impairment of bumblebee natural foraging behaviour induced by sublethal pesticide exposure. Functional Ecology, 28, 1459–1471.

Goulson, D., Nicholls, E., Botias, C. & Rotheray, E.L. (2015) Bee declines driven by combined stress from parasites, pesticides, and lack of flowers. Science, 347, 1255957.

Gradish, A.E., Steen, J.v.d., Scott-Dupree, C.D., Cabrera, A.R., Cutler, G.C., Goulson, D., Klein, O., Lehmann, D.M., Lückmann, J., O’Neill, B., Raine, N.E., Sharma, B. & Thompson, H. (2018) Comparison of Pesticide Exposure in Honey Bees (Hymenoptera: Apidae) and Bumble Bees (Hymenoptera: Apidae): Implications for Risk Assessments. Environmental Entomology, 48, 12–21, 10.

Hahn, D.A. & Denlinger, D.L. (2007) Meeting the energetic demands of insect diapause: nutrient storage and utilization. J Insect Physiol, 53, 760–773.

Heimpel, G.E. & de Boer, J.G. (2008) Sex determination in the hymenoptera. Annu Rev Entomol, 53, 209–230.

Jeschke, P., Nauen, R., Schindler, M. & Elbert, A. (2011) Overview of the status and global strategy for neonicotinoids. Journal of agricultural and food chemistry, 59, 2897–2908.

Jordan, A., Patch, H.M., Grozinger, C.M. & Khanna, V. (2021) Economic dependence and vulnerability of United States agricultural sector on insect-mediated pollination service. Environmental science & technology, 55, 2243–2253.

Keeling, C.I., Plettner, E. & Slessor, K.N. (2004) Hymenopteran semiochemicals. Top Curr Chem, 239, 133–177.

Klein, A.M., Vaissiere, B.E., Cane, J.H., Steffan-Dewenter, I., Cunningham, S.A., Kremen, C. & Tscharntke, T. (2007) Importance of pollinators in changing landscapes for world crops. Proc Biol Sci, 274, 303–313.

Kozii, I.V., Barnsley, S., Silva, M.C.B.d., Wood, S.C., Klein, C.D., de Mattos, I.M., Zabrodski, M.W., Silva, R.d.C.M., Fabela, C.I.O., Guillemin, L., Dvylyuk, I., Ferrari, M.C.O. & Simko, E. (2021) Reproductive fitness of honey bee queens exposed to thiamethoxam during development. Veterinary Pathology, 58, 1107–1118.

Krieger, G.M., Duchateau, M.J., Van Doorn, A., Ibarra, F., Francke, W. & Ayasse, M. (2006) Identification of queen sex pheromone components of the bumblebee *Bombus terrestris*. J Chem Ecol, 32, 453–471.

Krischik, V.A., Landmark, A.L. & Heimpel, G.E. (2007) Soil-Applied Imidacloprid Is Translocated to Nectar and Kills Nectar-Feeding *Anagyrus pseudococci* (Girault) (Hymenoptera: Encyrtidae). Environmental Entomology, 36, 1238–1245.

Krupke, C.H., Hunt, G.J., Eitzer, B.D., Andino, G. & Given, K. (2012) Multiple routes of pesticide exposure for honey bees living near agricultural fields. PLoS One, 7, e29268.

Kubo, R., Harano, K.-i. & Ono, M. (2017) Male scent-marking pheromone of *Bombus ardens ardens* (Hymenoptera; Apidae) attracts both conspecific queens and males. The Science of Nature, 104, 1–5.

Laycock, I., Lenthall, K.M., Barratt, A.T. & Cresswell, J.E. (2012) Effects of imidacloprid, a neonicotinoid pesticide, on reproduction in worker bumble bees (*Bombus terrestris*). Ecotoxicology, 21, 1937–1945.

Lecocq, T., Coppée, A., Mathy, T., Lhomme, P., Cammaerts-Tricot, M.-C., Urbanová, K., Valterová, I. & Rasmont, P. (2015) Subspecific differentiation in male reproductive traits and virgin queen preferences, in *Bombus terrestris*. Apidologie, 46, 595–605.

Li, S., Yu, X. & Feng, Q. (2019) Fat Body Biology in the Last Decade. Annual review of entomology, 64, 315–333.

Lürling, M. & Scheffer, M. (2007) Info-disruption: pollution and the transfer of chemical information between organisms. Trends in Ecology & Evolution, 22, 374–379.

Luxová, A., Urbanová, K., Valterová, I., Terzo, M. & Borg-Karlson, A.K. (2004) Absolute configuration of chiral terpenes in marking pheromones of bumblebees and cuckoo bumblebees. *Chirality: The Pharmacological*, Biological, and Chemical Consequences of Molecular Asymmetry, 16, 228–233.

Main, A.R., Webb, E.B., Goyne, K.W. & Mengel, D. (2020) Reduced species richness of native bees in field margins associated with neonicotinoid concentrations in non-target soils. Agriculture, Ecosystems & Environment, 287, 106693.

Minnameyer, A., Strobl, V., Bruckner, S., Camenzind, D.W., Van Oystaeyen, A., Wäckers, F., Williams, G.R., Yañez, O., Neumann, P. & Straub, L. (2021) Eusocial insect declines: Insecticide impairs sperm and feeding glands in bumblebees. Science of The Total Environment, 785, 146955.

Moffat, C., Buckland, S.T., Samson, A.J., McArthur, R., Chamosa Pino, V., Bollan, K.A., Huang, J.T.-J. & Connolly, C.N. (2016) Neonicotinoids target distinct nicotinic acetylcholine receptors and neurons, leading to differential risks to bumblebees. Scientific reports, 6, 1–10.

Müller, C. (2018) Impacts of sublethal insecticide exposure on insects — Facts and knowledge gaps. Basic and Applied Ecology, 30, 1–10.

Muth, F., Francis, J.S. & Leonard, A.S. (2019) Modality-specific impairment of learning by a neonicotinoid pesticide. Biology letters, 15, 20190359.

Navarro-Roldán, M.A. & Gemeno, C. (2017) Sublethal Effects of Neonicotinoid Insecticide on Calling Behavior and Pheromone Production of Tortricid Moths. Journal of Chemical Ecology, 43, 881–890.

Ollerton, J., Winfree, R. & Tarrant, S. (2011) How many flowering plants are pollinated by animals? Oikos, 120, 321–326.

Plowright, R. & Laverty, T. (1984) The ecology and sociobiology of bumble bees. Annual review of entomology, 29, 175–199.

Potts, S.G., Biesmeijer, J.C., Kremen, C., Neumann, P., Schweiger, O. & Kunin, W.E. (2010) Global pollinator declines: trends, impacts and drivers. Trends Ecol Evol, 25, 345–353.

Potts, S.G., Imperatriz-Fonseca, V., Ngo, H.T., Aizen, M.A., Biesmeijer, J.C., Breeze, T.D., Dicks, L.V., Garibaldi, L.A., Hill, R. & Settele, J. (2016) Safeguarding pollinators and their values to human well-being. Nature, 540, 220–229.

Raine, N.E. & Rundlöf, M. (2024) Pesticide Exposure and Effects on Non-Apis Bees. Annual review of entomology, 69, 551–576.

Ridley, M. (1990) The control and frequency of mating in insects. Functional Ecology, 75–84.

Rondeau, S., Baert, N., McArt, S. & Raine, N.E. (2022) Quantifying exposure of bumblebee (*Bombus spp.*) queens to pesticide residues when hibernating in agricultural soils. Environmental Pollution, 309, 119722.

Rundlöf, M., Andersson, G.K., Bommarco, R., Fries, I., Hederström, V., Herbertsson, L., Jonsson, O., Klatt, B.K., Pedersen, T.R. & Yourstone, J. (2015) Seed coating with a neonicotinoid insecticide negatively affects wild bees. Nature, 521, 77–80.

Sánchez-Bayo, F. & Wyckhuys, K.A.G. (2019) Worldwide decline of the entomofauna: A review of its drivers. Biological Conservation, 232, 8–27.

Sandrock, C., Tanadini, L.G., Pettis, J.S., Biesmeijer, J.C., Potts, S.G. & Neumann, P. (2014) Sublethal neonicotinoid insecticide exposure reduces solitary bee reproductive success. Agricultural and Forest Entomology, 16, 119–128.

Schmid-Hempel, R. & Schmid-Hempel, P. (2000) Female mating frequencies in *Bombus spp*. from Central Europe. Insectes Sociaux, 47, 36–41.

Schmuck, R., Schöning, R., Stork, A. & Schramel, O. (2001) Risk posed to honeybees (*Apis mellifera* L, Hymenoptera) by an imidacloprid seed dressing of sunflowers. Pest Management Science: formerly Pesticide Science, 57, 225–238.

Sgolastra, F., Hinarejos, S., Pitts-Singer, T.L., Boyle, N.K., Joseph, T., Lūckmann, J., Raine, N.E., Singh, R., Williams, N.M. & Bosch, J. (2018) Pesticide Exposure Assessment Paradigm for Solitary Bees. Environmental Entomology, 48, 22–35.

Siviter, H., Horner, J., Brown, M.J. & Leadbeater, E. (2020) Sulfoxaflor exposure reduces egg laying in bumblebees *Bombus terrestris*. Journal of Applied Ecology, 57, 160–169.

Siviter, H. & Muth, F. (2022) Exposure to the novel insecticide flupyradifurone impairs bumblebee feeding motivation, learning, and memory retention. Environmental Pollution, 307, 119575.

Smith, J.M. & Harper, D. (2003) Animal signals. Oxford University Press.

Stanley, D.A., Smith, K.E. & Raine, N.E. (2015) Bumblebee learning and memory is impaired by chronic exposure to a neonicotinoid pesticide. Sci Rep, 5, 16508.

Starr, C.K. (1984) Sperm competition, kinship, and sociality in the aculeate Hymenoptera. Sperm competition and the evolution of animal mating systems, 428, 459.

Stoner, K.A. & Eitzer, B.D. (2012) Movement of soil-applied imidacloprid and thiamethoxam into nectar and pollen of squash (Cucurbita pepo). PLoS One, 7, e39114.

Straub, L., Villamar-Bouza, L., Bruckner, S., Chantawannakul, P., Gauthier, L., Khongphinitbunjong, K., Retschnig, G., Troxler, A., Vidondo, B., Neumann, P. & Williams, G.R. (2016) Neonicotinoid insecticides can serve as inadvertent insect contraceptives. Proc Biol Sci, 283.

Stuligross, C. & Williams, N.M. (2020) Pesticide and resource stressors additively impair wild bee reproduction. Proceedings of the Royal Society B, 287, 20201390.

Suchail, S., De Sousa, G., Rahmani, R. & Belzunces, L.P. (2004) In vivo distribution and metabolisation of 14C-imidacloprid in different compartments of *Apis mellifera* L. Pest Management Science: formerly Pesticide Science, 60, 1056–1062.

Suchail, S., Debrauwer, L. & Belzunces, L.P. (2004) Metabolism of imidacloprid in *Apis mellifera*. Pest Management Science: formerly Pesticide Science, 60, 291–296.

Svensson, B. & Bergstrom, G. (1977) Volatile marking secretions from the labial gland of North European Pyrobombus D.T. males (Hymenoptera, Apidae).

Tappert, L., Pokorny, T., Hofferberth, J. & Ruther, J. (2017) Sublethal doses of imidacloprid disrupt sexual communication and host finding in a parasitoid wasp. Scientific reports, 7, 1–9.

Tasman, K., Rands, S.A. & Hodge, J.J.L. (2020) The Neonicotinoid Insecticide Imidacloprid Disrupts Bumblebee Foraging Rhythms and Sleep. iScience, 23, 101827.

Tosi, S., Burgio, G. & Nieh, J.C. (2017) A common neonicotinoid pesticide, thiamethoxam, impairs honey bee flight ability. Sci Rep, 7, 1201.

Tosi, S., Sfeir, C., Carnesecchi, E., vanEngelsdorp, D. & Chauzat, M.-P. (2022) Lethal, sublethal, and combined effects of pesticides on bees: A meta-analysis and new risk assessment tools. Science of The Total Environment, 844, 156857.

Treanore, E. & Amsalem, E. (2020) The effect of intrinsic physiological traits on diapause survival and their underlying mechanisms in an annual bee species *Bombus impatiens*. Conservation Physiology, 8, coaa103.

Treanore, E., Barie, K., Derstine, N., Gadebusch, K., Orlova, M., Porter, M., Purnell, F. & Amsalem, E. (2021) Optimizing Laboratory Rearing of a Key Pollinator, *Bombus impatiens*. Insects, 12, 673.

Tricoire-Leignel, H., Thany, S., Gadenne, C. & Anton, S. (2012) Pest Insect Olfaction in an Insecticide-Contaminated Environment: Info-Disruption or Hormesis Effect. Frontiers in Physiology, 3.

Tsvetkov, N., Samson-Robert, O., Sood, K., Patel, H., Malena, D., Gajiwala, P., Maciukiewicz, P., Fournier, V. & Zayed, A. (2017) Chronic exposure to neonicotinoids reduces honey bee health near corn crops. Science, 356, 1395–1397.

Urbanová, K., Cahlíková, L., Hovorka, O., Ptáček, V. & Valterová, I. (2008) Age-dependent changes in the chemistry of exocrine glands of *Bombus terrestris* queens. Journal of Chemical Ecology, 34, 458–466.

Valterová, I., Martinet, B., Michez, D., Rasmont, P. & Brasero, N. (2019) Sexual attraction: a review of bumblebee male pheromones. Z Naturforsch C J Biosci, 74, 233–250.

van Honk, C.G.J., Velthuis, H.H.W. & Röseler, P.F. (1978) A sex pheromone from the mandibular glands in bumblebee queens. Experientia, 34, 838–839.

Van Wilgenburg, E., Driessen, G. & Beukeboom, L.W. (2006) Single locus complementary sex determination in Hymenoptera: an “unintelligent” design? Frontiers in Zoology, 3, 1–15.

Whitehorn, P.R., O’Connor, S., Wackers, F.L. & Goulson, D. (2012) Neonicotinoid pesticide reduces bumble bee colony growth and queen production. Science, 336, 351–352.

Whitehorn, P.R., Tinsley, M.C. & Goulson, D. (2009) Kin recognition and inbreeding reluctance in bumblebees. Apidologie, 40, 627–633.

Willis Chan, D.S. & Raine, N.E. (2021) Population decline in a ground-nesting solitary squash bee (*Eucera pruinosa*) following exposure to a neonicotinoid insecticide treated crop (*Cucurbita pepo*). Scientific reports, 11, 4241.

Wu-Smart, J. & Spivak, M. (2018) Effects of neonicotinoid imidacloprid exposure on bumble bee (Hymenoptera: Apidae) queen survival and nest initiation. Environmental Entomology, 47, 55–62.

Yew, J.Y. & Chung, H. (2015) Insect pheromones: An overview of function, form, and discovery. Progress in Lipid Research, 59, 88–105.

Zioga, E., Kelly, R., White, B. & Stout, J.C. (2020) Plant protection product residues in plant pollen and nectar: A review of current knowledge. Environmental Research, 189, 109873.

