## Supplementary Material for "Sublethal pesticide exposure decreases mating and disrupts chemical signaling in a beneficial pollinator"

Table S1 – Statistical results of pairwise comparisons of imidacloprid amount in treatment groups.

| Comparison | Z | P.unadj | P.adj |
|---|---|---|---|
| Gyne 60ppb - Gyne 6ppb | 2.397 | 0.017 | **0.021** |
| Gyne 60ppb - Control | 4.791 | <0.001 | **<0.001** |
| Gyne 6ppb - Control | 2.458 | 0.014 | **0.028** |
| Gyne 60ppb - Male 60ppb | -0.662 | 0.508 | 0.564 |
| Gyne 6ppb - Male 60ppb | -2.995 | 0.003 | **0.007** |
| Control - Male 60ppb | -5.315 | <0.001 | **<0.001** |
| Gyne 60ppb - Male 6ppb | 2.405 | 0.016 | **0.023** |
| Gyne 6ppb - Male 6ppb | 0.008 | 0.994 | 0.994 |
| Control - Male 6ppb | -2.450 | 0.014 | **0.024** |
| Male 60ppb - Male 6ppb | 3.003 | 0.003 | **0.009** |

Table S2 – The relative abundance (percent ± standard error) of compounds included in analyses of gyne cuticular hydrocarbons for the treatment groups control (0 ppb), 6 ppb, and 60 ppb imidacloprid. Compound names are abbreviated such that 9-$C_{23}$ indicates 9-tricosene, etc. RI is the linear retention index.

| **Compound** | **RI** | **Control % ± SE** | **6 ppb % ± SE** | **60 ppb % ± SE** |
|---|---|---|---|---|
| $C_{21}$ | 2100 | 0.54 ± 0.07 | 0.44 ± 0.04 | 0.45 ± 0.03 |
| 9-$C_{23}$ | 2270 | 2.55 ± 0.15 | 2.3 ± 0.1 | 3.02 ± 0.1 |
| $C_{23}$ | 2300 | 8.76 ± 0.81 | 8.39 ± 0.29 | 9.2 ± 0.4 |
| 11-$C_{25}$ | 2467 | 0.61 ± 0.07 | 0.54 ± 0.08 | 0.43 ± 0.03 |
| 9-$C_{25}$ | 2474 | 15.33 ± 0.46 | 15.65 ± 0.59 | 18.06 ± 0.5 |
| 7-$C_{25}$ | 2479 | 1.7 ± 0.15 | 1.59 ± 0.08 | 1.81 ± 0.11 |
| $C_{25}$ | 2500 | 11.95 ± 0.31 | 11 ± 0.25 | 11.37 ± 0.27 |
| $C_{26}$ | 2600 | 0.56 ± 0.02 | 0.54 ± 0.02 | 0.53 ± 0.01 |
| 9-$C_{27}$ | 2674 | 9.92 ± 0.39 | 10.55 ± 0.25 | 10.42 ± 0.24 |
| 7-$C_{27}$ | 2680 | 3.7 ± 0.15 | 3.79 ± 0.16 | 3.61 ± 0.18 |
| $C_{27}$ | 2700 | 12.02 ± 0.46 | 11.79 ± 0.43 | 10.98 ± 0.31 |
| 9-$C_{29}$ | 2875 | 8.19 ± 0.39 | 8.97 ± 0.18 | 8.06 ± 0.16 |
| 7-$C_{29}$ | 2881 | 2.01 ± 0.08 | 2.07 ± 0.09 | 1.74 ± 0.06 |
| $C_{29}$ | 2900 | 2.2 ± 0.1 | 2.21 ± 0.11 | 1.89 ± 0.08 |
| 9-$C_{31}$ | 3074 | 3.12 ± 0.15 | 3.66 ± 0.08 | 3.05 ± 0.08 |
| UNK_06 | 3912 | 1.9 ± 0.22 | 2.11 ± 0.11 | 2.16 ± 0.12 |
| UNK_07 | 3927 | 0.73 ± 0.08 | 0.83 ± 0.06 | 0.97 ± 0.07 |
| UNK_08 | 3940 | 0.94 ± 0.1 | 0.85 ± 0.07 | 0.53 ± 0.04 |

Table S3 – The relative abundance (percent ± standard error) of compounds included in analyses of male cuticular hydrocarbons for the treatment groups control (0 ppb), 6 ppb, and 60 ppb imidacloprid. Compound names are abbreviated such that 9-$C_{25}$ indicates 9-pentacosene, etc. RI is the linear retention index.

| **Compound** | **RI** | **Control % ± SE** | **6 ppb % ± SE** | **60 ppb % ± SE** |
|---|---|---|---|---|
| $C_{23}$ | 2300 | 7.35 ± 0.34 | 6.88 ± 0.35 | 7.27 ± 0.28 |
| 9-$C_{25}$ | 2474 | 4.6 ± 0.23 | 4.56 ± 0.22 | 5.12 ± 0.25 |
| 7-$C_{25}$ | 2482 | 1.4 ± 0.06 | 1.37 ± 0.06 | 1.55 ± 0.06 |
| $C_{25}$ | 2500 | 21.58 ± 0.35 | 21.01 ± 0.31 | 20.15 ± 0.32 |
| $C_{26}$ | 2599 | 1.31 ± 0.04 | 1.24 ± 0.05 | 1.03 ± 0.07 |
| 9-$C_{27}$ | 2676 | 5.81 ± 0.19 | 5.59 ± 0.18 | 6.47 ± 0.17 |
| 7-$C_{27}$ | 2684 | 6.52 ± 0.21 | 6.53 ± 0.19 | 7.22 ± 0.24 |
| 5-$C_{27}$ | 2694 | 0.78 ± 0.03 | 0.79 ± 0.03 | 0.91 ± 0.03 |
| $C_{27}$ | 2700 | 26.65 ± 0.61 | 26.86 ± 0.59 | 24.85 ± 0.56 |
| 9-$C_{29}$ | 2876 | 6.05 ± 0.19 | 5.92 ± 0.15 | 6.61 ± 0.15 |
| 7-$C_{29}$ | 2884 | 4.78 ± 0.15 | 4.99 ± 0.11 | 5.32 ± 0.16 |
| $C_{29}$ | 2900 | 6.9 ± 0.29 | 7.57 ± 0.21 | 6.73 ± 0.29 |
| 9-$C_{31}$ | 3078 | 4.65 ± 0.2 | 4.86 ± 0.13 | 5.14 ± 0.17 |
| 7-$C_{31}$ | 3086 | 0.77 ± 0.04 | 0.85 ± 0.05 | 0.84 ± 0.05 |
| $C_{31}$ | 3100 | 0.87 ± 0.06 | 0.98 ± 0.05 | 0.81 ± 0.05 |

Table S4 – The relative abundance (percent ± standard error) of compounds included in analyses of gyne mandibular glands for the treatment groups control (0 ppb), 6 ppb, and 60 ppb imidacloprid. Some compound names are abbreviated such that 9-$C_{23}$ indicates 9-tricosene. As the extract was derivatized with BSTFA, compounds with carboxyl or hydroxyl groups have trimethylsilyl adducts (TMS). RI is the linear retention index.

| **Compound** | **RI** | **Control Percent ± SE** | **6 ppb Percent ± SE** | **60 ppb Percent ± SE** |
|---:|---:|---:|---:|---:|
| phosphoric acid - triTMS | 1289 | 7.93 ± 1.06 | 10.23 ± 1.21 | 10 ± 0.9 |
| 3-hydroxy hexanoic acid - diTMS | 1318 | 1.35 ± 0.33 | 1.38 ± 0.35 | 1.6 ± 0.34 |
| 3-hydroxy octanoic acid - diTMS | 1490 | 15.43 ± 2.51 | 13.18 ± 1.52 | 16.21 ± 2.6 |
| unknown_01 | 1559 | 19.75 ± 3.02 | 22.81 ± 4.32 | 23.1 ± 5.74 |
| 3-hydroxy decanoic acid - diTMS | 1668 | 13.04 ± 2.18 | 9.76 ± 1.28 | 12.94 ± 2.07 |
| 3-hydroxy dodecanoic acid - diTMS | 1850 | 1.9 ± 0.35 | 1.53 ± 0.28 | 1.94 ± 0.39 |
| unknown_02 | 1915 | 0.21 ± 0.05 | 0.55 ± 0.16 | 0.45 ± 0.2 |
| unknown_03 | 1926 | 0.36 ± 0.14 | 0.65 ± 0.14 | 0.53 ± 0.14 |
| hexadecenoic acid I - TMS | 2026 | 0.91 ± 0.37 | 1.23 ± 0.31 | 0.96 ± 0.24 |
| hexadecenoic acid II - TMS | 2038 | 4.14 ± 1.65 | 6.01 ± 1.74 | 4.36 ± 1.61 |
| hexadecanoic acid - TMS | 2046 | 1.2 ± 0.3 | 1.34 ± 0.27 | 1.34 ± 0.29 |
| octadecenoic acid I - TMS | 2217 | 7.3 ± 2.14 | 9.27 ± 1.77 | 7.23 ± 2.06 |
| octadecenoic acid II - TMS | 2222 | 1.01 ± 0.33 | 1.47 ± 0.32 | 1.23 ± 0.4 |
| octadecenoic acid - TMS | 2241 | 1.02 ± 0.26 | 1.73 ± 0.36 | 1.53 ± 0.39 |
| 9-$C_{23}$ | 2269 | 1.17 ± 0.24 | 0.72 ± 0.15 | 1.02 ± 0.27 |
| $C_{23}$ | 2300 | 1.61 ± 0.44 | 1.05 ± 0.27 | 1.07 ± 0.31 |
| 9-$C_{25}$ | 2470 | 6.66 ± 1.46 | 4.27 ± 1.04 | 4.24 ± 1.06 |
| 7-$C_{25}$ | 2477 | 0.96 ± 0.23 | 1 ± 0.27 | 0.84 ± 0.21 |
| $C_{25}$ | 2500 | 1.41 ± 0.29 | 1.02 ± 0.21 | 0.93 ± 0.18 |
| 9-$C_{27}$ | 2671 | 2.72 ± 0.57 | 2 ± 0.52 | 1.78 ± 0.36 |
| 7-$C_{27}$ | 2679 | 1.47 ± 0.37 | 1.34 ± 0.4 | 0.96 ± 0.2 |
| $C_{27}$ | 2700 | 1.19 ± 0.29 | 1.09 ± 0.3 | 0.79 ± 0.15 |
| 9-$C_{29}$ | 2871 | 1.66 ± 0.33 | 1.2 ± 0.27 | 1.04 ± 0.17 |
| 7-$C_{29}$ | 2879 | 1.03 ± 0.25 | 0.66 ± 0.17 | 0.48 ± 0.08 |
| $C_{29}$ | 2900 | 0.45 ± 0.11 | 0.32 ± 0.08 | 0.2 ± 0.03 |
| 9-$C_{31}$ | 3072 | 0.99 ± 0.2 | 0.77 ± 0.15 | 0.58 ± 0.1 |
| unknown_04 | 3266 | 1.42 ± 0.28 | 1.53 ± 0.23 | 1.35 ± 0.27 |
| unknown_05 | 3535 | 0.89 ± 0.25 | 1.09 ± 0.48 | 0.58 ± 0.14 |
| unknown_06 | 3738 | 0.83 ± 0.25 | 0.79 ± 0.25 | 0.71 ± 0.23 |

Table S5 – The relative abundance (percent ± standard error) of compounds included in analyses of male labial glands for the treatment groups control (0 ppb), 6 ppb, and 60 ppb imidacloprid. Compound names are abbreviated such that $C_{23}$ indicates tricosane. As the extract was derivatized with BSTFA, compounds with carboxyl or hydroxyl groups have trimethylsilyl adducts (TMS). RI is the linear retention index.

| Compound | RI | Control Percent ± SE | 6 ppb Percent ± SE | 60 ppb Percent ± SE |
|---|---|---|---|---|
| 2,3-dihydrofarnesol-TMS | 1768 | 18.66 ± 0.41 | 18.21 ± 0.73 | 18.5 ± 0.97 |
| hexadecenoic acid I - TMS | 2035 | 57.53 ± 1.46 | 59.03 ± 1.13 | 56.77 ± 1.65 |
| unknown 1 | 2220 | 1.2 ± 0.07 | 1.22 ± 0.08 | 1.27 ± 0.09 |
| unknown 2 | 2227 | 1.21 ± 0.06 | 1.17 ± 0.04 | 1.23 ± 0.06 |
| unknown 3 | 2246 | 0.97 ± 0.04 | 0.95 ± 0.03 | 0.97 ± 0.06 |
| $C_{23}$ | 2300 | 2.94 ± 0.26 | 2.74 ± 0.18 | 3.03 ± 0.36 |
| $C_{25}$ | 2500 | 2.19 ± 0.18 | 2.04 ± 0.13 | 2.21 ± 0.19 |
| $C_{27}$ | 2700 | 1.37 ± 0.12 | 1.26 ± 0.08 | 1.39 ± 0.12 |
| unknown 4 | 2751 | 1.37 ± 0.07 | 1.23 ± 0.06 | 1.4 ± 0.11 |
| farnesyl ester I | 3148 | 6.05 ± 0.48 | 5.94 ± 0.34 | 6.51 ± 0.73 |
| farnesyl ester V | 3347 | 1.48 ± 0.16 | 1.32 ± 0.09 | 1.46 ± 0.16 |

Table S6 – GC-MS fragment ions from dimethyl disulfide derivatization used to identify double bond positions in cuticular alkenes.

| **Compound** | **(M)$^+$** | **(M-47)$^+$** | **(A)$^+$** | **(B)$^+$** |
|---|---|---|---|---|
| 9-$C_{23}$ | 416 | 369 | 173 | 243 |
| 11-$C_{25}$ | 444 | 397 | 201 | 243 |
| 9-$C_{25}$ | 444 | 397 | 173 | 271 |
| 7-$C_{25}$ | 444 | 397 | 145 | 299 |
| 9-$C_{27}$ | 472 | 425 | 173 | 299 |
| 7-$C_{27}$ | 472 | 425 | 145 | 327 |
| 5-$C_{27}$ | 472 | 425 | 117 | 355 |
| 9-$C_{29}$ | 500 | 453 | 173 | 327 |
| 7-$C_{29}$ | 500 | 453 | 145 | 355 |
| 9-$C_{31}$ | 528 | 481 | 173 | 355 |

Figure S1- Example GC-MS spectrum of DMDS derivatized 9-tricosene (9-$C_{23}$) showing diagnostic ions. Labeled fragments correspond to those described in Table S6 (following (Carlson, Roan, Yost, & Hector, 2002).

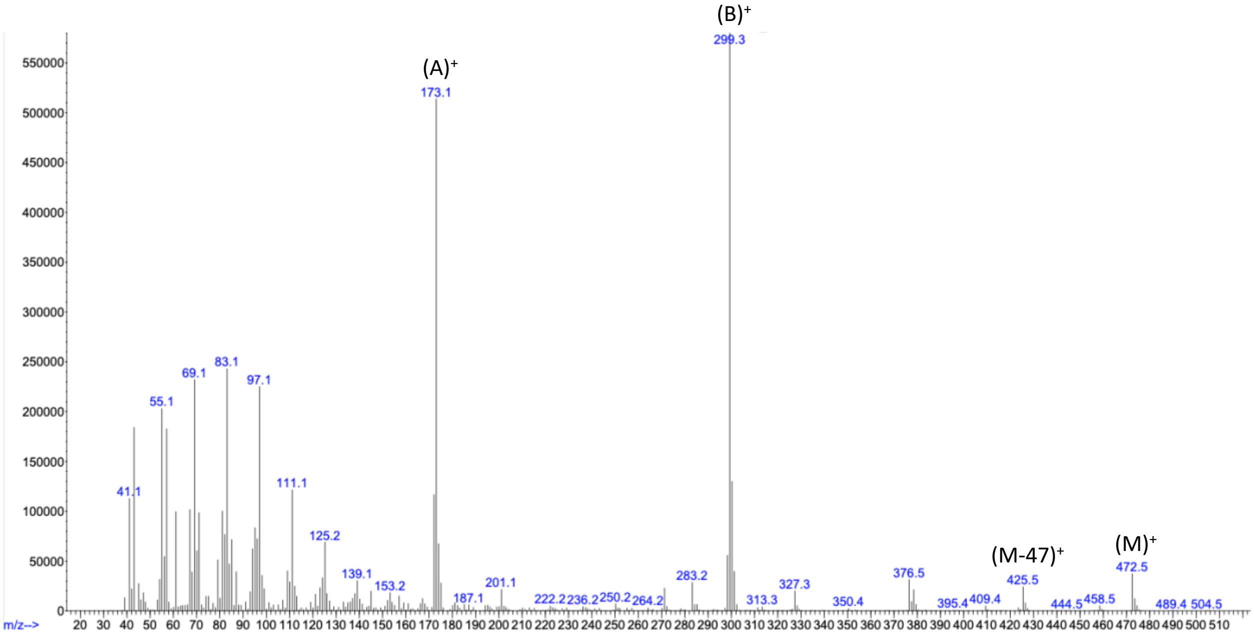

Figure S2 – Size differences between all gynes and males measured in the study. Panel A shows wet weight (grams) and panel B shows head width (mm).

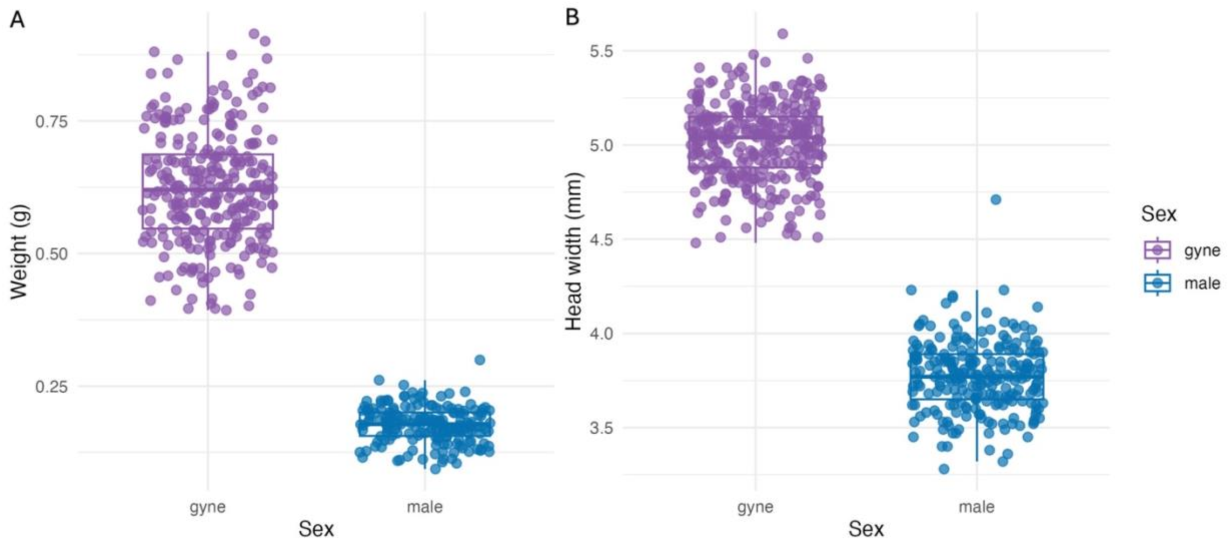

References

Carlson, D. A., Roan, C. S., Yost, R. A., & Hector, J. (2002). Dimethyl disulfide derivatives of long chain alkenes, alkadienes, and alkatrienes for gas chromatography/mass spectrometry. *Analytical Chemistry*, *61*(14), 1564-1571. doi:10.1021/ac00189a019
